# A mitochondrial surveillance mechanism activated by *SRSF2* mutations in hematologic malignancies

**DOI:** 10.1101/2023.06.25.546449

**Authors:** Xiaolei Liu, Sudhish A. Devadiga, Robert F. Stanley, Ryan Morrow, Kevin Janssen, Mathieu Quesnel-Vallières, Oz Pomp, Adam A. Moverley, Chenchen Li, Nicolas Skuli, Martin P. Carroll, Jian Huang, Douglas C. Wallace, Kristen W. Lynch, Omar Abdel-Wahab, Peter S. Klein

## Abstract

Splicing factor mutations are common in myelodysplastic syndrome (MDS) and acute myeloid leukemia (AML), but how they alter cellular functions is unclear. We show that the pathogenic *SRSF2^P95H/+^* mutation disrupts the splicing of mitochondrial mRNAs, impairs mitochondrial complex I function, and robustly increases mitophagy. We also identified a mitochondrial surveillance mechanism by which mitochondrial dysfunction modifies splicing of the mitophagy activator *PINK1* to remove a poison intron, increasing the stability and abundance of *PINK1* mRNA and protein. *SRSF2^P95H^*-induced mitochondrial dysfunction increased *PINK1* expression through this mechanism, which is essential for survival of *SRSF2^P95H/+^* cells. Inhibition of splicing with a glycogen synthase kinase 3 inhibitor promoted retention of the poison intron, impairing mitophagy and activating apoptosis in *SRSF2^P95H/+^* cells. These data reveal a homeostatic mechanism for sensing mitochondrial stress through *PINK1* splicing and identify increased mitophagy as a disease marker and a therapeutic vulnerability in *SRSF2^P95H^* mutant MDS and AML.

## INTRODUCTION

Myelodysplastic syndrome (MDS) is a clonal hematopoietic disorder characterized by hyperproliferative bone marrow, dysplastic hematopoietic cells, peripheral cytopenias, and high risk of progression to acute myeloid leukemia (AML). Recurrent mutations in RNA splicing factors (SFs), including *SF3B1*, *SRSF2, and U2AF1*, account for 60-70% of somatic mutations in MDS (1) and are common in myeloproliferative neoplasms (MPNs), the MDS/MPN overlap disorder chonic myelomonocytic leukemia (CMML), and AML (1–6). However, the mechanisms by which these mutations alter cellular functions or contribute to transformation remain unclear.

Mutations in distinct SFs cause myeloid disorders with similar phenotypic features but limited overlap in specific mRNAs that are aberrantly spliced. This may suggest that distinct mRNA targets are responsible for these myeloid malignancies or that shared pathways are affected downstream of aberrantly spliced mRNAs. Hotspot mutations in *SRSF2* alter RNA binding site specificity and disrupt the splicing of mRNAs encoding hematopoietic regulators that are frequently mutated in AML (7, 8). For example, the *SRSF2^P95*^* mutation promotes inclusion of a poison exon in *EZH2* mRNA, leading to nonsense mediated decay and reduced levels of *EZH2* (7). Thus, the number of patients with reduced *EZH2* expression is higher than the number with *EZH2* mutations (8). These studies support a pathogenic mechanism targeting the splicing of a single mRNA, but do not rule out additional contributions from other splicing events. Furthermore, transcriptomic analyses of different SF mutants show that altered splicing of distinct mRNAs can affect common downstream pathways, including splicing itself, protein synthesis, and mitochondrial function (9).

Splicing factor mutations occur as mutually exclusive, heterozygous point mutations at specific residues (10, 11). The presence of a wild-type splicing factor is required for survival of MDS cells that express a mutant splicing factor, suggesting vulnerability associated with disrupted splicing factor activity. Consistent with this, hematopoietic cells with heterozygous splicing factor mutations are sensitized to additional perturbation of the splicing machinery by small molecule splicing inhibitors (12, 13). This observation has led to clinical trials with splicing inhibitors in refractory leukemia patients (12, 14–18). However, these trials have been ineffective so far (14), and there are currently no FDA-approved therapies that target splicing factors in MDS or AML.

Glycogen synthase kinase-3α/β (GSK-3α/β) are multifunctional serine/threonine kinases encoded by two similar genes, *GSK3A* and *GSK3B*. GSK-3 functions downstream of Flt3, c-Kit, and Wnt signaling, all pathways associated with MDS and AML (19). *Gsk3a/b* knockdown (20) or *Gsk3b* deletion (21) in mouse bone marrow causes a pronounced myeloproliferative phenotype and *Gsk3a/b* double knockout (DKO) causes marked expansion of both mature granulocytes and primitive blasts, leading to an aggressive AML (21), but the substrates that mediate GSK-3 functions in mouse and human hematopoietic malignancies have not been extensively characterized. Our recent phosphoproteomic analysis of GSK-3 substrates identified multiple core splicing factors implicated in MDS and AML (19, 22), consistent with prior studies supporting the importance of GSK-3 in splicing regulation, including evidence that GSK-3 phosphorylation of SRSF2 regulates splicing of *MAPT* (Tau) exon 10 (23–25). Both pharmacological inhibition and genetic loss of *GSK3* disrupt pre-mRNA splicing on a transcriptome-wide scale in diverse cell types (19, 26), raising the possibility that clinically well tolerated GSK-3 inhibitors could be repurposed to treat splicing factor mutant MDS and AML.

In the present study, we assessed transcriptome-wide changes in splicing upon GSK-3 inhibition (GSK-3i) in leukemic cells. We find that GSK-3i disrupts splicing and selectively kills MDS and AML cells with SF mutations. Mechanistically, GSK-3i induces a heightened splicing response in SF mutant cells by repressing cassette exon inclusion and promoting intron retention. These findings led us to focus on SRSF2, as *SRSF2* mutant cells show disrupted splicing of nuclear-encoded mitochondrial genes, impaired mitochondrial function, and an increase in mitophagy that is essential for their survival. Splicing of mRNA encoding the mitochondrial surveillance factor *PINK1* is altered in response to mitochondrial stress in *SRSF2^P95H/+^* mutant cells, leading to a more stable splice form and reflecting what we believe to be a novel mechanism for sensing mitochondrial stress. GSK-3i disrupts splicing of *PINK1*, promoting a splice form with a premature stop codon and reducing overall *PINK1* mRNA levels. The reduction in *PINK1* is associated with reduced mitophagy and enhanced cell death in *SRSF2^P95H/+^* mutant cells, sparing cells with wild-type *SRSF2*. Our findings reveal a mechanism for mitochondrial surveillance and identify a new therapeutic vulnerability in *SRSF2* mutant MDS and AML.

## RESULTS

### Preferential sensitivity of spliceosomal mutant leukemia to GSK-3i

Heterozygous splicing factor mutations sensitize leukemic cells to additional perturbation of the core splicing machinery (13). As GSK-3 regulates splicing in other cell types (22–26), we tested whether GSK-3i would enhance cell death in leukemic cells with heterozygous knockins of recurrent driver mutations in *SRSF2* and *SF3B1*. Isogenic *SRSF2^P95H/+^* and *SF3B1^K700E/+^* knock-in K562 cells (13) were cultured with the GSK-3 inhibitor CHIR99021 (CHIR) or vehicle control and proliferation was monitored at 2 and 4 days. Both *SRSF2^P95H/+^* and *SF3B1^K700E/+^* cells were preferentially sensitive to CHIR (Figure 1A and Supplemental Figure 1A) and the clinically well tolerated GSK-3 inhibitor lithium (Supplemental Figure 1B) compared to isogenic cells with wild-type (*WT*) *SRSF2* and *SF3B1*. A flow cytometric assay for apoptosis revealed that GSK-3i promotes both early and late apoptosis (Figure 1B) in splicing factor mutant cells compared to parental cells but had little impact on their cell cycle status (Supplemental Figure 1C). Collectively, these data demonstrate that GSK-3i preferentially kills spliceosomal mutant leukemias.

**Figure 1.**
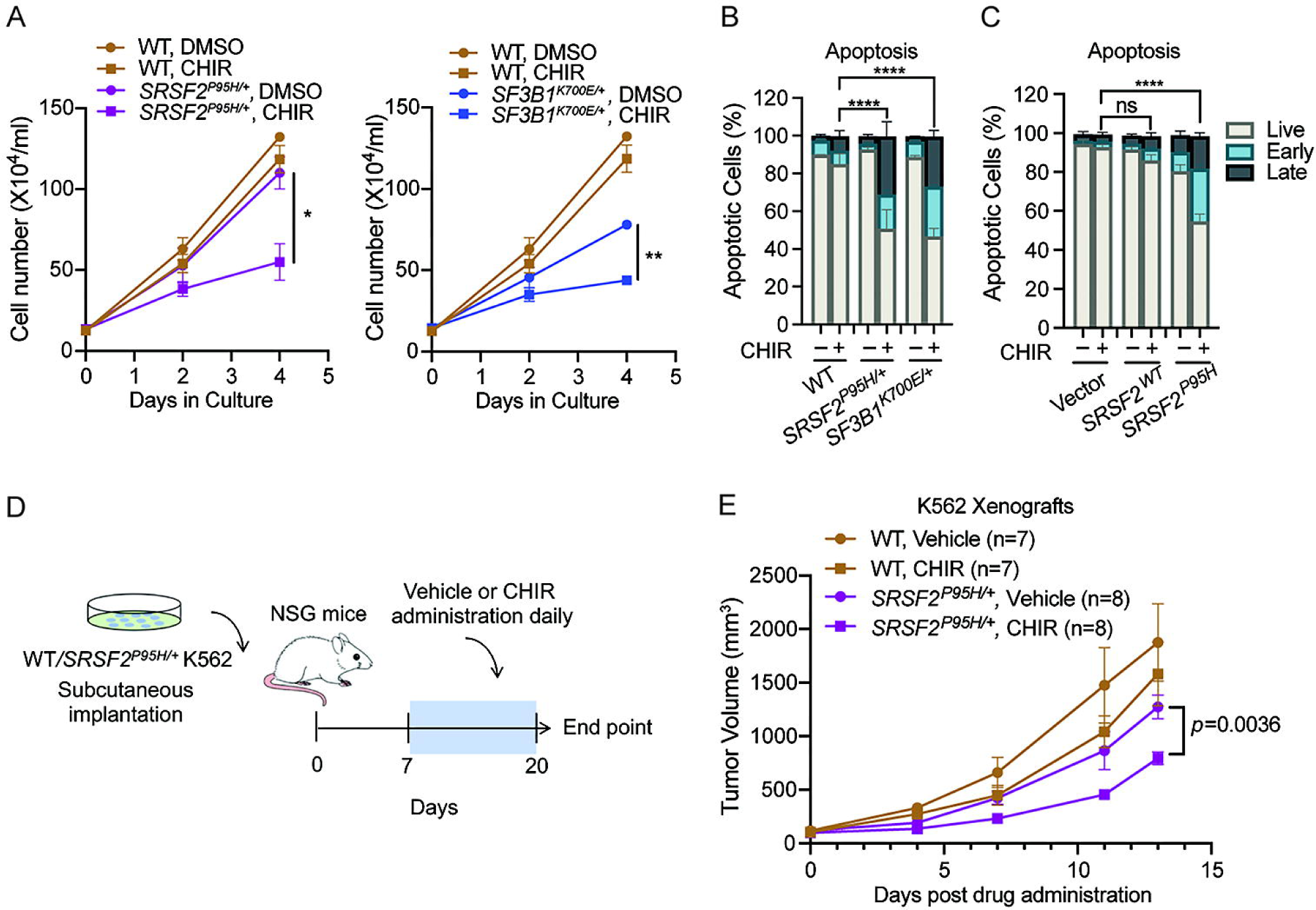
Preferential cytotoxicity of GSK-3i in SF-mutant leukemia over WT counterparts. (**A**) *WT, SRSF2^P95H/+^*and *SF3B1^K700E/+^* cells were cultured with DMSO or 3uM CHIR, and cell numbers were counted at 2 d and 4 d (mean ± SD). (**B**) Percentages of viable, early-, and late-apoptotic cells based on 7-AAD and Annexin V flow cytometric analysis of isogenic K562 *WT*, *SRSF2^P95H/+^* and *SF3B1^K700E/+^* cells treated with DMSO or 3uM CHIR *in vitro* for 4 d (mean ± SD). Statistical analysis of viable fraction in mutant cells treated with CHIR relative to *WT* counterparts was performed using a two-tailed Chi-squared test. **** *p* < 0.0001. (**C**) Percentages of viable, early-, and late-apoptotic cells of K562 parental, *SRSF2^wt^*-, or *SRSF2^P95H^*-overexprssing cells treated with vehicle or 3uM CHIR *in vitro* for 4 d (mean ± SD). Statistical analysis of viable fraction in *SRSF2^P95H^*-overexpressing cells treated with CHIR relative to *SRSF2^wt^* conterparts was performed using a two-tailed Chi-squared test. **** *p* < 0.0001. (**D**) Schematic of *in vivo* K562 xenograft experiment. (**E**) Mean tumor volume in NSG mice subcutaneously implanted with K562 isogenic *WT* or *SRSF2^P95H/+^*cells. Mice received subcutaneous injections of vehicle or CHIR (30mg/kg) daily. Mean tumor volumes ± SEM are shown. For data in A, and E, \**p* < 0.05 and \*\**p* < 0.01 (2-way ANOVA with Sidak’s multiple comparisons test).

As an alternative approach to compare the effect of mutated and wild-type *SRSF2* in otherwise genetically identical cells, we introduced *SRSF2^P95H^* or *SRSF2^wt^* into parental TF-1 or K562 cells along with mCherry (Supplemental Figure 1D, E), and then followed proliferation of mCherry positive and untransduced cells. Expression of *SRSF2^P95H^*resulted in a competitive disadvantage when compared with WT-expressing cells (Supplemental Figure 1F), consistent with prior reports (7, 27). Further, GSK-3i markedly increased apoptosis in TF-1 and K562 cells expressing *SRSF2^P95H^* compared to parental cells and cells overexpressing *SRSF2^wt^* (Figure 1C and Supplemental Figure 1G). Therefore, *SRSF2^P95H^* confers sensitivity to GSK-3i whether introduced as a heterozygous knockin or overexpressed in a genetically identical background. We next tested whether GSK-3i selectively kills splicing factor mutant cells *in vivo* in mice bearing xenografts of isogenic *WT* and *SRSF2^P95H/+^* K562 cells (Figure 1D). Daily administration of CHIR slowed the growth of xenografts with *SRSF2^P95H/+^* (*p*=0.0036) but had no significant effect on *SRSF2^wt^* xenografts (Figure 1E and Supplemental Figure 1H, I).

### GSK-3i induces cell death in splicing factor mutant cells from patients with hematologic malignancies

Based on the promising results with leukemic cell lines (Figure 1), we evaluated the preferential cytotoxicity of GSK-3i in primary cells from patients with AML or the MDS/MPN overlap disorder CMML. GSK-3i induced apoptosis in primary cells from patients with CMML or AML with *SRSF2*, *SF3B1*, or *U2AF1* mutations, but not in leukemic cells with wild-type splicing factors or in CD34^+^ cells from healthy subjects (Figure 2A, and Supplemental Table 1).

**Figure 2.**
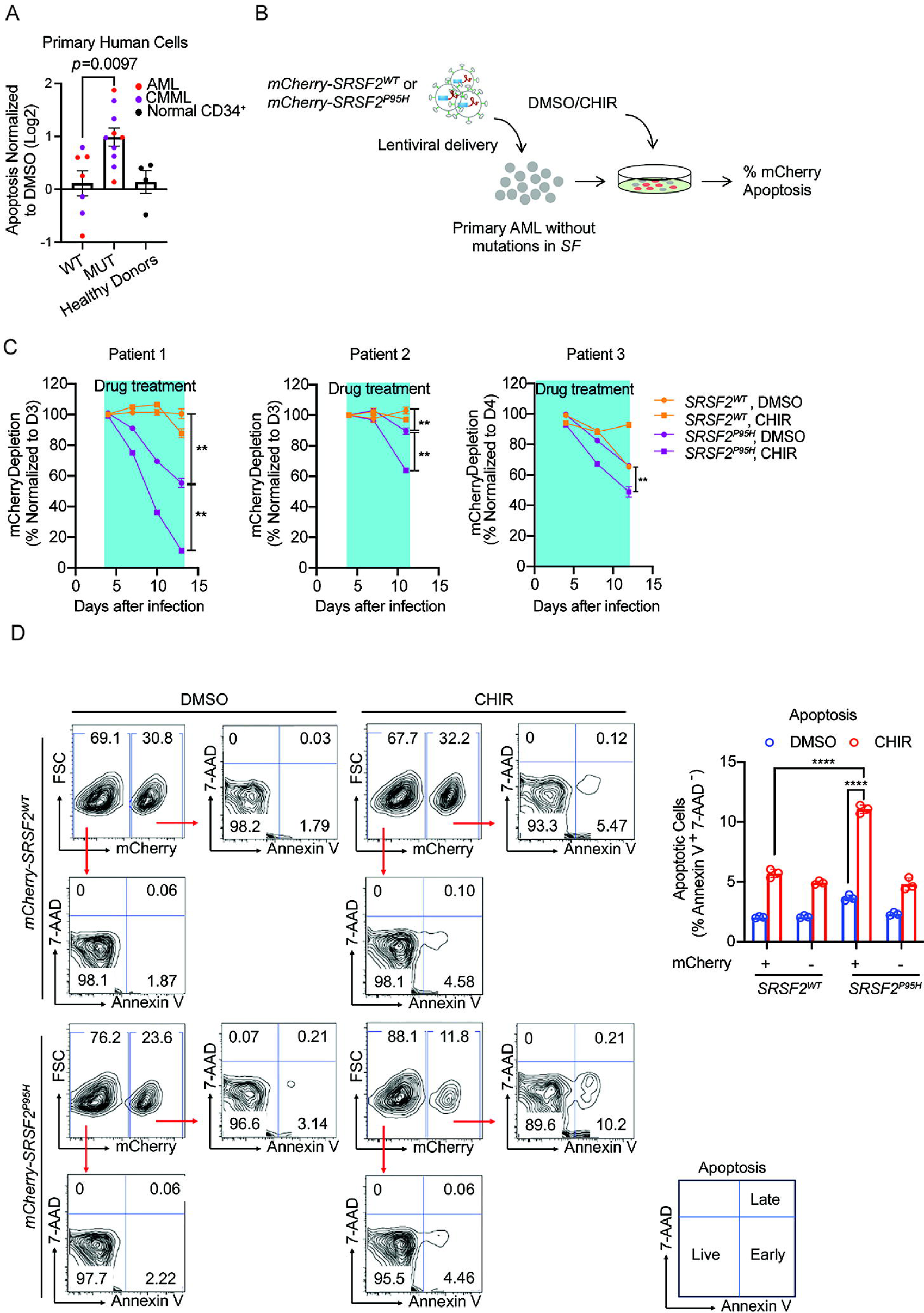
Selective cytotoxicity of GSK-3i in primary human leukemia cells with SF mutations. (**A**) CHIR induced higher levels of apoptosis in splicing factor mutant (mut) cells from patients with CMML (red) or AML (purple) compared to patients with *wt* splicing factors or CD34^+^ cells from healthy donors (black). Log2[fold change] in apoptosis in CHIR relative to DMSO treated is shown. Genetic information for each patient is shown in Data file S1. Statistical analysis was performed using a two-tailed Mann Whitney test. (**B**) Schematic of lentivirus infection in primary AML cells for apoptosis assay. (**C**) Primary AML cells from 3 patients were transduced with lentivirus encoding *WT* or mutant *SRSF2* as in panel B and cultured in the absence or presence of 3uM CHIR (blue box indicates duration of drug treatment). The percentage of mCherry^+^ cells normalized to the number at day 3 (patients 1 and 2) or day 4 (patient 3) is shown. (**D**) Representative flow cytometric analysis (left) and quantification (right) of apoptosis in primary cells from patients with AML overexpressing either *SRSF2^+^* or *SRSF2^P95H^*, as measured by Annexin V and 7-AAD staining in absence or presence of 3uM CHIR. Data in C and D are presented as mean ± SD; \*\**p* < 0.01 and \*\*\*\**p* < 0.0001 (2-way ANOVA with Sidak’s multiple comparisons test).

Given the heterogeneity of primary human AML cells, we also tested the preferential cytotoxicity of GSK-3i in an isogenic context by expressing *SRSF2^WT^* or *SRSF2^P95H^* with mCherry in primary blasts from AML patients with wild-type splicing factors and then assessing the loss of mCherry positive cells over time with or without CHIR treatment (Figure 2B and Supplemental Table 1). Primary AML cells expressing *SRSF2^P95H^* were preferentially depleted upon CHIR exposure (Figure 2C). In contrast, GSK-3i had no substantial effect on primary AML cells expressing *WT SRSF2* (Figure 2C). Cells expressing mutant *SRSF2* also displayed a higher level of apoptosis in response to CHIR treatment relative to controls transduced with wild-type *SRSF2* (Figure 2D and Supplemental Figure 2). These data confirm that *SRSF2^P95H^* confers sensitivity to GSK-3i in primary AML cells.

### GSK-3i alters global gene expression and splicing in human leukemic cells

To understand the functional consequences of GSK-3i on gene expression and splicing, we performed deep RNA-sequencing (RNA-seq, 10^8^ reads/replicate sample) of parental K562 and isogenic *SRSF2^P95H/+^*and *SF3B1^K700E/+^* lines treated with or without CHIR (Figure 3A and Supplemental Figure 3. CHIR treatment had distinct effects on the transcriptomes of WT, *SRSF2^P95H/+^*, and *SF3B1^K700E/+^* mutant cells (Supplemental Figure 3B, C, and Supplemental Table 2). Additionally, GSK-3i altered the expression of multiple *BCL2* family genes in a manner that may contribute to cell death, including decreased expression of the anti-apoptotic *BCL-XL* (*BCL2L1*-encoded) and increased expression of the pro-apoptotic genes *BAK1*, *BCL2L11*, and *BIK* regardless of SF mutation status (Supplemental Figure 3D).

**Figure 3.**
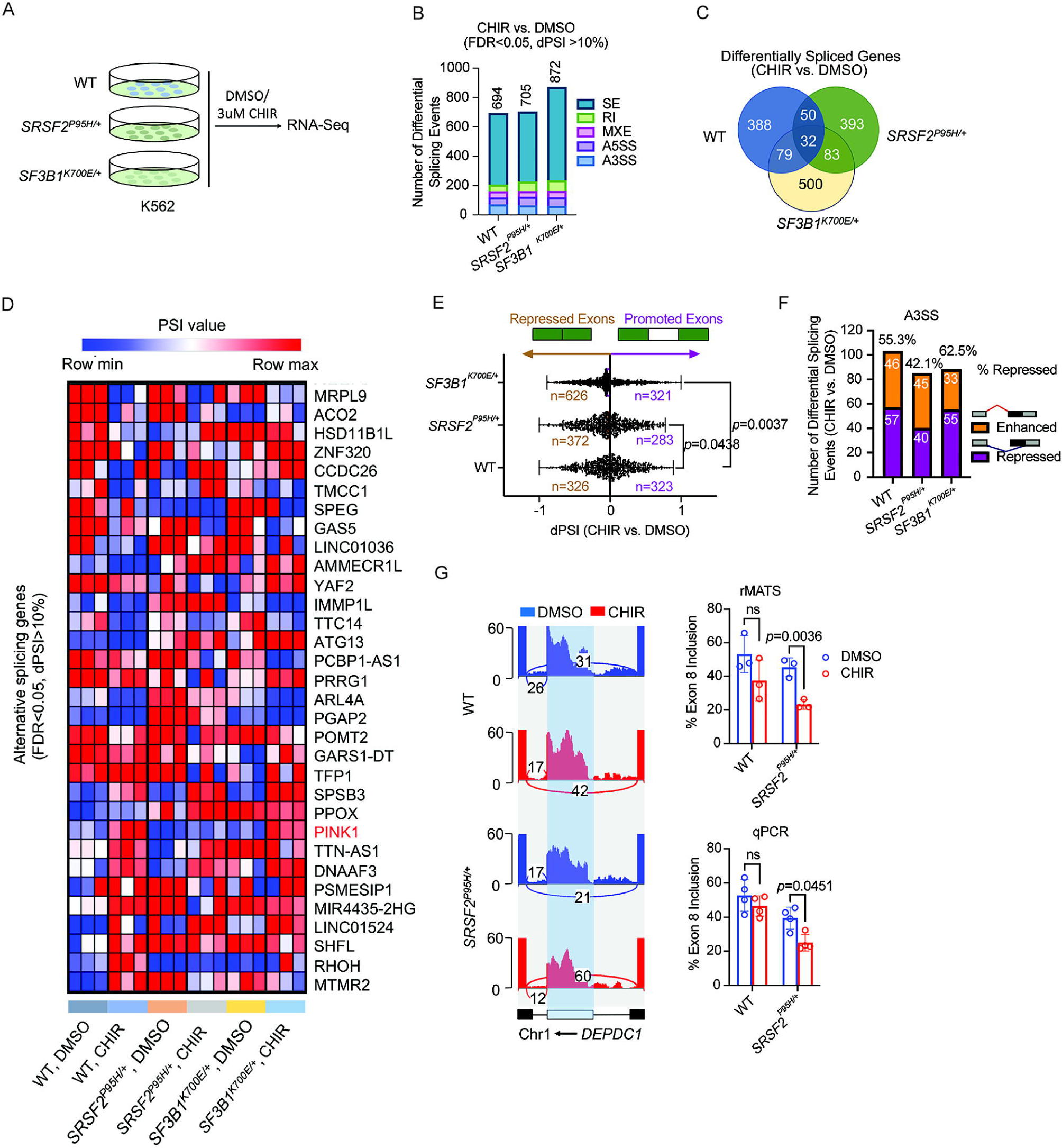
GSK-3i is associated with global alterations in gene expression and splicing in human leukemic cells. (**A**) Schematic of deep RNA-Seq in *WT*, *SRSF2^P95H/+^* and *SF3B1^K700E/+^* K562 cells treated with DMSO or 3uM CHIR for 24 h. (**B**) Bar graphs show numbers of differential splicing events (FDR<0.05, dPSI>10%) in *WT*, *SRSF2^P95H/+^* and *SF3B1^K700E/+^* K562 cells treated with CHIR versus DMSO controls. A3SS, alternative 3’ splice site; A5SS, alternative 5’ splice site; MXE, mutually exclusive exon; RI, retained intron; SE, skipped exon. (**C**) Venn diagram showing the number of overlapping alternatively spliced genes between CHIR and DMSO in *WT*, *SRSF2^P95H/+^* and *SF3B1^K700E/+^* K562 cells. (**D**) Heat map of PSI values for overlapping alternatively spliced genes comparing CHIR and DMSO in *WT*, *SRSF2^P95H/+^* and *SF3B1^K700E/+^* K562 cells. (**E**) Scatterplots of cassette exon inclusion in *WT*, *SRSF2^P95H/+^* and *SF3B1^K700E/+^* K562 cells treated with 3uM CHIR relative to DMSO treated controls. Numbers in brown (left) and purple (right) indicate number of cassette exons whose inclusion is repressed or promoted, respectively, in CHIR treated relative to DMSO treated cells. *p* values were determined by 1-way ANOVA with Sidak’s multiple comparisons test. (**F**) Bar graphs show numbers of alternative 3’ splice site events upon CHIR treatment compared to DMSO in *WT*, *SRSF2^P95H/+^* and *SF3B1^K700E/+^* K562 cells. Purple and orange indicate intron-proximal 3’ splice sites whose usage is repressed or enhanced, respectively, in CHIR treated relative to DMSO treated cells. (**G**) Sashimi plots of *DEPDC1* in *WT* and *SRSF2^P95H/+^* cells treated with DMSO or CHIR (left). Bar plots shows quantification of percentage of exon inclusion based on rMATS analysis (top right) and on isoform specific qPCR validation (bottom right). Percentage of exon inclusion was quantified by RT-qPCR analysis of mRNA levels containing the cassette exons normalized to total mRNA levels. Data are presented as the mean ± SD. *p* values were determined by 2-way ANOVA with Sidak’s multiple comparisons test. ns: not significant.

We then performed genome-wide analysis of splicing using the rMATS pipeline (28). We measured the change in percentage spliced in (dPSI) values across five main types of alternative splicing (AS) events (skipped exon [SE], alternative 5’ splice-site exon [A5SS], alternative 3’ ss exon [A3SS], mutually exclusive exons [MXE], and retained introns [RI]) in CHIR-treated versus control (Supplemental Table 3). Using a false discovery rate (FDR) < 0.05 and dPSI >10% (Figure 3B), we identified 694 alternative splicing events in parental cells, 705 in *SRSF2^P95H/+^* and 872 in *SF3B1^K700E/+^* cells. However, only 32 differentially spliced mRNAs were found in all the three genotypes upon GSK-3i, suggesting that GSK-3i has distinct and independent consequences on splicing in *WT* and splicing factor mutant cells (Figure 3C and 3D). The predominant change in RNA splicing upon CHIR treatment in *SRSF2^P95H/+^* and *SF3B1^K700E/+^*cells was increased cassette exon skipping relative to DMSO-treat controls, while approximately equal proportions of exon inclusion and exclusion with CHIR exposure were observed in parental cells (Figure 3E). SF3B1 normally facilitates 3′ splice site recognition and is involved in the splicing of most introns by binding to the branchpoint (29). GSK-3i in *SF3B1^K700E/+^*cells resulted in a higher proportion of repressed A3’SS (Figure 3F) and increased number of RI events (Figure 3B). Consistent with the distinct effects of GSK-3i on splicing in cells with *WT* vs mutant SFs, multiple regulators of hematopoietic cell survival and proliferation were alternatively spliced only in SF mutant K562 cells (Figure 3G and Supplemental Figure 3E). Among them, GSK-3i enhanced exon skipping in *DEPDC1* pre-mRNA in *SRSF2^P95H/+^* cells compared to wild-type counterparts, an event that was further validated by transcript specific RT-qPCR (Figure 3G). Taken together, inhibition of GSK-3 further reduces splicing fidelity in splicing factor mutant hematopoietic cells, likely explaining the selective toxicity of GSK-3i to splicing factor mutant leukemias over wild-type counterparts.

### GSK-3 regulates *PINK1* splicing

To test whether genes that are alternatively spliced following GSK-3i contribute to the preferential killing of spliceosome-mutant cells, we evaluated transcripts that exhibit concomitant dysregulation in gene expression and splicing with CHIR treatment (Figure 4A). We focused our attention initially on *PINK1* because it is a serine/threonine kinase that serves as a critical sensor of mitochondrial damage and activates mitophagy to maintain mitochondrial homeostasis (30). GSK-3i reduces the abundance and alters the splicing of *PINK1* mRNA in *WT*, *SRSF2^P95H/+^*, and *SF3B1^K700E/+^*K562 cells. A Sashimi plot shows that GSK-3i enhances the retention of intron-6, which includes a premature termination codon (PTC) that is predicted to promote mRNA degradation through nonsense mediated decay (NMD) (Figure 4B). RT-qPCR confirms that GSK-3i reduces the abundance of *PINK1* in *WT*, *SRSF2^P95H/+^*, and *SF3B1^K700E/+^* cells (Figure 4C). This change in *PINK1* abundance was not due to a change in transcription, as detection of primary transcripts with PCR primers that detect intron-1 or intron-5 showed no significant difference in nascent transcripts with GSK-3i in *WT* or splicing factor mutant cells (Figure 4D). Qualitative RT-PCR with primers that span exons 6 and 7 confirmed that GSK-3 inhibition with CHIR (Figure 4E) or AR-A014418 (Supplemental Figure 4) impaired excision of intron-6, increasing intron-6 retention (568bp), in *SRSF2^+/+^*, *SRSF2^P95H/+^*, and *SF3B1^K700E/+^*K562 cells. CHIR also promoted retention of intron-6 in primary cells from patients with AML, CMML, and CD34^+^ cells from healthy donors (Figure 4F). To confirm that CHIR impairs *PINK1* splicing through inhibition of GSK-3, we tested *PINK1* splicing in *WT* and *GSK3A/B* DKO cells. Similar to CHIR and AR-A014418, *GSK3A/B* DKO impaired intron-6 excision. In contrast, overexpression of *GSK3B* in DKO cells restored excision of intron-6 (Figure 4G). Blocking NMD by knockdown of *UPF1*, a core NMD factor, increased the mRNA stability and expression of the intron 6-retained form of *PINK1* (Figure 4H). Taken together, these data show that GSK-3 regulates *PINK1* splicing and that CHIR reduces *PINK1* mRNA abundance specifically by inhibiting GSK-3 and promoting retention of an intron containing an in-frame PTC.

**Figure 4.**
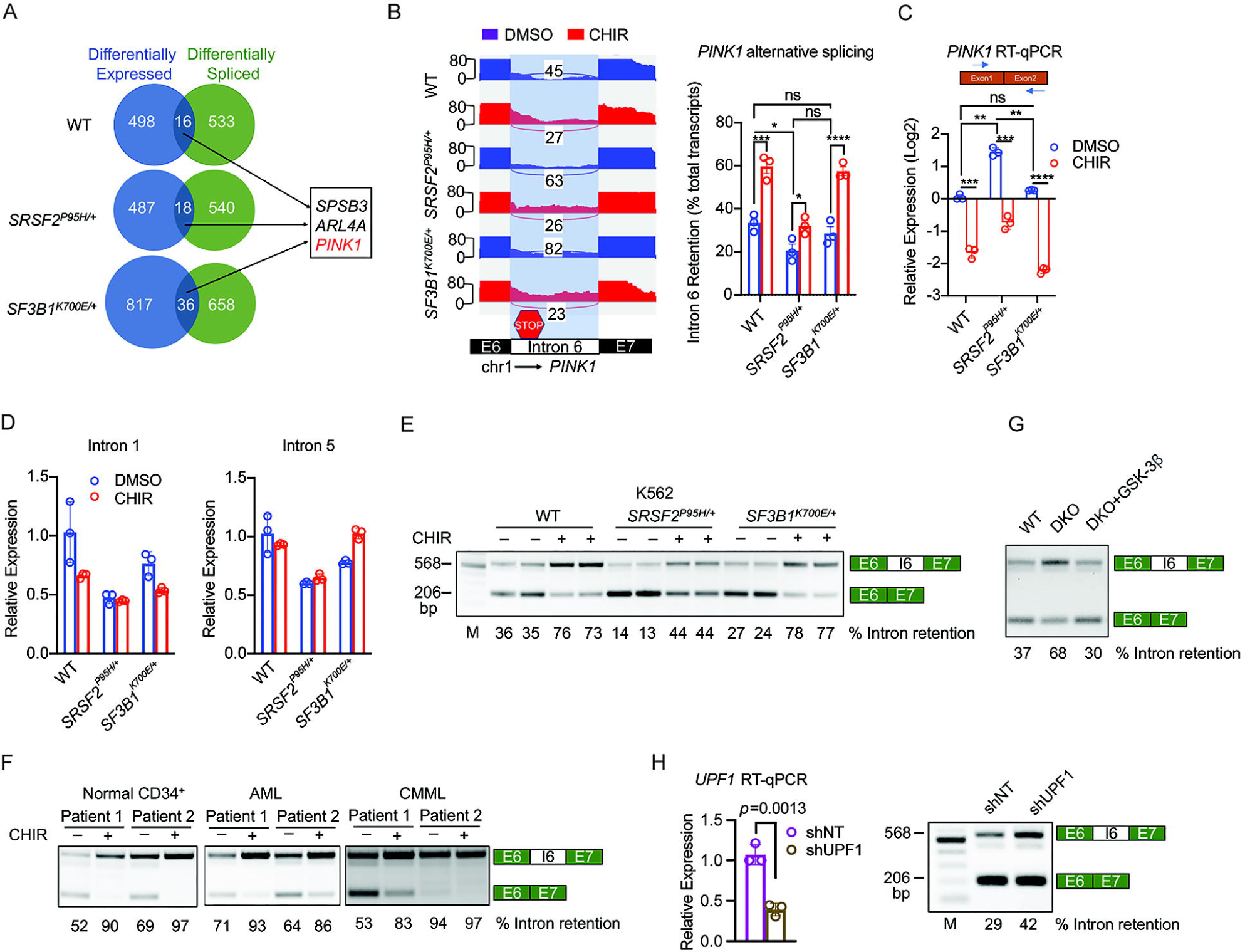
GSK-3 regulates *PINK1* splicing. (**A**) Venn diagram of genes differentially expressed (blue) or differentially spliced (green) in CHIR-treated *WT*, *SRSF2^P95H/+^* and *SF3B1^K700E/+^* K562 cells relative to DMSO-treated controls. (**B**) Sashimi plots of *PINK1* in *WT*, *SRSF2^P95H/+^*and *SF3B1^K700E/+^* cells treated with DMSO or CHIR (left). Bar graph shows intron 6 retention as a percentage of total transcripts (right). (**C**) *PINK1* mRNA levels in *WT*, *SRSF2^P95H/+^* and *SF3B1^K700E/+^* cells treated with DMSO or 3uM CHIR for 24 h detected by RT-qPCR using primers that span exons 1 and 2 (mean ± SD). For data in B and C, * *p* < 0.05, ** *p* < 0.01, *** *p* < 0.001, and \*\*\*\**p*<0.0001 (2-way ANOVA with Sidak’s multiple comparisons test). (**D**) *PINK1* nascent transcript detection by RT-qPCR using primers for intron 1 (left) and intron 5 (right) in *WT*, *SRSF2^P95H/+^* and *SF3B1^K700E/+^* cells treated with DMSO or 3uM CHIR for 24 h. (**E**-**F**) Representative RT-PCR with primers spanning exon 6 (E6), intron 6 (I6), and exon 7 (E7) of *PINK1* in *WT*, *SRSF2^P95H/+^*, *SF3B1^K700E/+^* K562 cells (E) and in CD34^+^ cells from healthy donors, primary AML cells, and CD34^+^ cells from CMML patients (F) treated with DMSO or 3uM CHIR for 24h. M: DNA marker. PCR product with retained intron 6 is 568bp and product for spliced exon6-7 (without retained intron) is 206bp. (**G**) RT-PCR analysis of *PINK1* splicing in *WT*, *GSK3A/B* DKO, and *GSK3A/B* DKO cells overexpressing GSK-3β (HEK293T). (**H**) *UPF1* mRNA levels detected in K562 cells transduced with lentivirus expressing shRNA against *UPF1* compared to non-targeting control (left), and levels of intron-retained and -spliced *PINK1* mRNAs following *UPF1* knockdown compared to control measured by RT-PCR (right). Data are presented as mean ± SD*. p* value was determined by Student’s *t* test.

### *SRSF2^P95H^* increases *PINK1* expression and mitophagy

In contrast to GSK-3i, the *SRSF2^P95H^* mutation markedly increases *PINK1* mRNA abundance compared to parental and *SF3B1^K700E/+^*cells (Figure 4C). Nascent *PINK1* transcripts were not increased in *SRSF2^P95H/+^*cells (Figure 4D), indicating that the increase in mRNA abundance was due to stabilization of the *PINK1* transcript rather than increased transcription. In support of this, excision of intron-6 was increased in *SRSF2^P95H/+^* cells compared to parental cells or *SF3B1^K700E/+^*cells (Figure 4E). Increased excision of intron-6 was further validated in TF-1 cells overexpressing *SRSF2^P95H^* compared to cells expressing *SRSF2^wt^*(Supplemental Figure 5A).

PINK1 activates mitophagy, and in parallel with the increase in *PINK1* levels, expression of other mitophagy-related genes, including *OPTN*, *ULK1*, *TOMM7*, *CALCOCO2*, *NBR1*, and *TAX1BP1,* was significantly increased in *SRSF2^P95H/+^* cells (Figure 5A). The increase in *PINK1* and other mitophagy markers was associated with a substantial increase in mitophagy specifically in *SRSF2^P95H/+^* cells, as determined by the colocalization of TOMM20 (mitochondrial marker) and LAMP1 (lysosomal marker) (Figure 5B). Immunofluorescent staining demonstrated that PINK1 colocalizes with TOMM20 (Supplemental Figure 5B) and PARKIN (Supplemental Figure 5C), suggesting that PINK1 was stabilized on impaired mitochondria in *SRSF2^P95H/+^* cells. Transmission electron microscopy (TEM) showed accumulation of autophagic vacuoles with defective mitochondria with swollen matrix and collapsed cristae enclosed by a double membrane in *SRSF2^P95H/+^* cells (Figure 5C). As *SRSF2^P95H^* and GSK-3i appear to have opposing effects on *PINK1* splicing and abundance, we tested the effect of CHIR on mitophagy in *SRSF2^P95H/+^* cells and found that GSK-3i impairs mitophagy in *SRSF2^P95H/+^* cells (Figure 5B and 5C), in parallel with the reduction in *PINK1* mRNA. We then assessed mitophagic flux by evaluating the accumulation of mitochondria with mito-tracker green (MTG) staining in the presence of chloroquine (CQ), which blocks the final step of autophagy/mitophagy, fusion with the lysosome, by inhibiting lysosomal acidification. CQ increased mitochondrial mass, as expected (Supplemental Figure 5D). The increase was significantly higher in *SRSF2^P95H/+^*cells than in *SRSF2^+/+^* cells. Mitophagic flux was then determined by subtracting the MTG value for untreated cells from the value for cells treated with CQ (31). The *SRSF2^P95H/+^*mutation significantly enhanced mitophagic flux (Figure 5D). A similar increase in mitophagic flux was observed with Lys05, an alternative inhibitor of lysosomal acidification (Figure 5D). GSK-3i also increased mitochondrial mass in both *SRSF2^+/+^* and *SRSF2^P95H/+^* cells, with or without CQ or Lys05 (Supplemental Figure 5D and 5E), consistent with reduced *PINK1* levels and impaired mitophagy. Western blot analysis confirmed that PINK1 protein is increased in *SRSF2^P95H/+^* cells and decreased upon CHIR treatment (Figure 5E).

**Figure 5.**
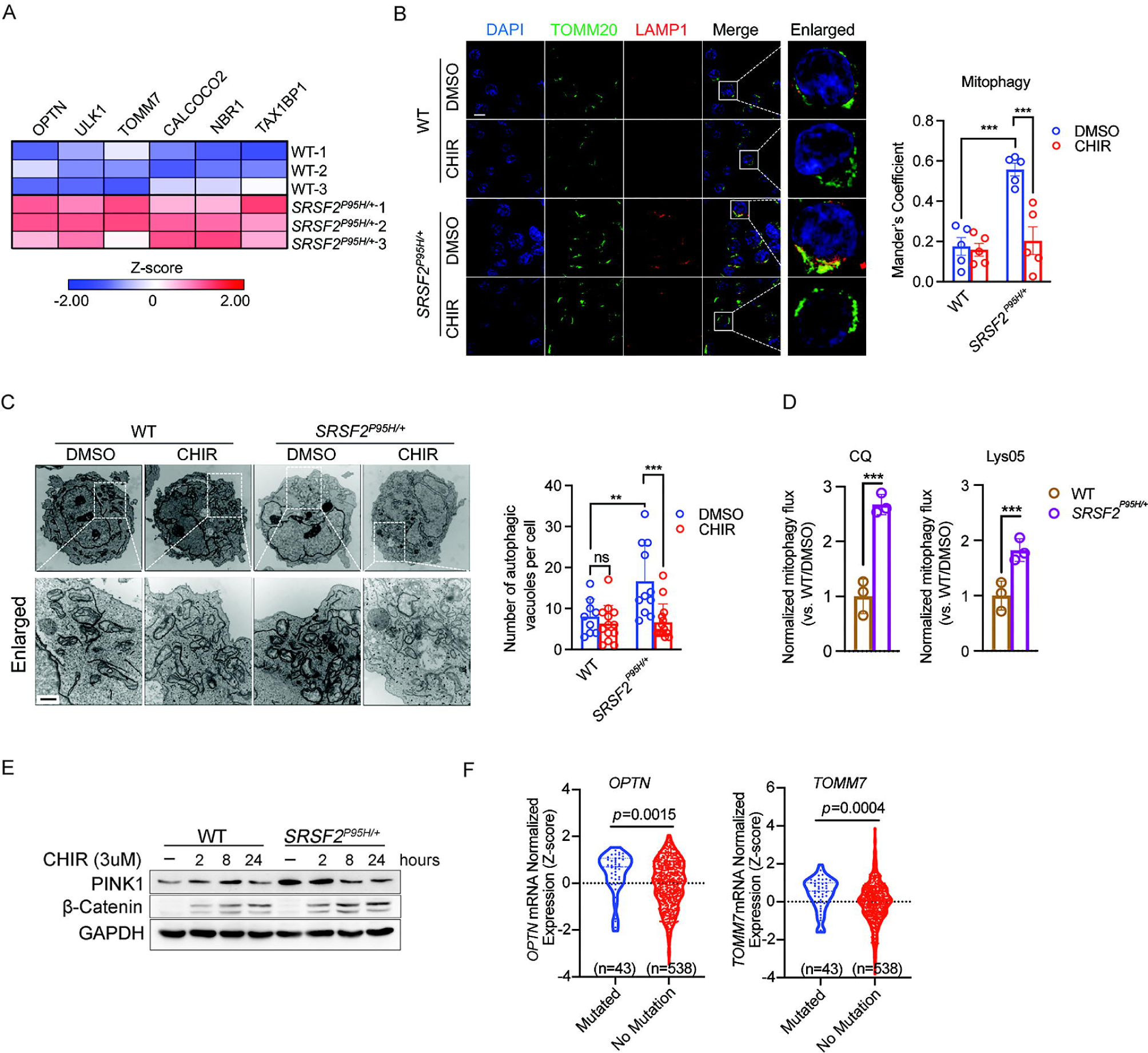
*SRSF2^P95H^* increases *PINK1* expression and mitophagy. (**A**) Heat map shows z-scored expression of mitophagy related genes in *WT* and *SRSF2^P95H/+^* K562 cells. (**B**) Representative confocal images of mitochondria (green, TOMM20^+^) and lysosomes (red, LAMP1^+^) in K562 *WT*, or *SRSF2^P95H/+^*cells treated with DMSO or 3uM CHIR for 2d (left). Scale bar, 10 μm. Bar plot shows quantification of the co-localization of mitochondria with lysosomes (n=5 fields for each group. N=3). (**C**) Representative TEM images of *WT* and *SRSF2^P95H/+^* K562 cells treated with DMSO or 3µM CHIR for 2d (left). Scale bar, 0.5 μm. Bar plot (right) shows quantification of the number of autophagic vacuoles per cell. Each circle represents one cell. (**D**) Bar graph shows mitophagic flux determined by mito-tracker green (MTG) staining in *WT* and *SRSF2^P95H/+^*K562 treated with or without 2uM CHIR for 48h. Mitochondrial net flux was calculated by mitochondrial accumulation in the presence of 100uM Chloroquine (CQ) or 50uM Lys05 for 4h (31). Data are presented as the mean ± SD. * *p*<0.05, ** *p*<0.01, *** *p*<0.001, **** *p*<0.0001, by 2-way ANOVA with Sidak’s multiple comparisons test (**B**-**D**). (**E**) Immunoblotting of PINK1 and β-catenin in *WT* and *SRSF2^P95H/+^* cells treated with DMSO or 3 uM CHIR for indicated times. (**F**) Violin plot of *OPTN*, and *TOMM7* normalized expression in AML patients in the TCGA dataset (n = 581) with or without mutations in *SRSF2*. Statistical analysis was performed using two-tailed Mann-Whitney test.

To address whether an increase in mitophagy is a feature of primary cells from patients with hematological malignancies, we queried the Cancer Genome Atlas database for expression of markers of mitophagy in hematologic malignancies with or without the *SRSF2* mutation. Expression of the canonical mitophagy markers *OPTN (p = 0.0015)*, and *TOMM7* (*p* = 0.0004) was significantly increased in patients with *SRSF2^P95*^* (n = 43) compared to patients with wild-type *SRSF2* (n = 538) (Figure 5F). No significant difference was observed between patients with *WT* and mutant *SF3B1* (Supplemental Figure 5F). These data show elevated mitophagy marker expression specifically in *SRSF2* mutant MDS and AML, and we therefore focused on the impact of *SRSF2^P95H^* on mitochondrial function and the role of GSK-3-dependent *PINK1* splicing in the regulation of mitophagy in *SRSF2^P95H/+^* cells.

### The *SRSF2* mutation is associated with accumulation of defective mitochondria

This increase in mitophagy indicates that the *SRSF2^P95H/+^*mutation causes mitochondrial dysfunction, which could arise through mis-splicing of nuclear mRNAs encoding mitochondrial proteins. We analyzed splicing variations in *SRSF2^P95H/+^* cells compared to *SRSF2^+/+^* cells (Supplemental Figure 6A). Gene ontology (GO) analysis of differentially spliced genes in *SRSF2^P95H/+^* cells revealed enrichment of processes related to regulation of protein targeting to mitochondrion organization, mitochondrial genome maintenance, mitochondrial respiratory chain complex I assembly, and oxidative phosphorylation (Figure 6A, and Supplemental Table 4). We identified 12 alternatively spliced mRNAs that are associated with mitochondrial respiratory chain complex I (Figure 6A, and Supplemental Figure 6B). These findings are consistent with GO analyses in *SRSF2* mutant cells from primary MDS (9), CMML (Figure 6B) (7), and AML patients (Figure 6B) (7, 32), which similarly showed enrichment of mitochondrial genes in the set of alternatively spliced mRNAs.

**Figure 6.**
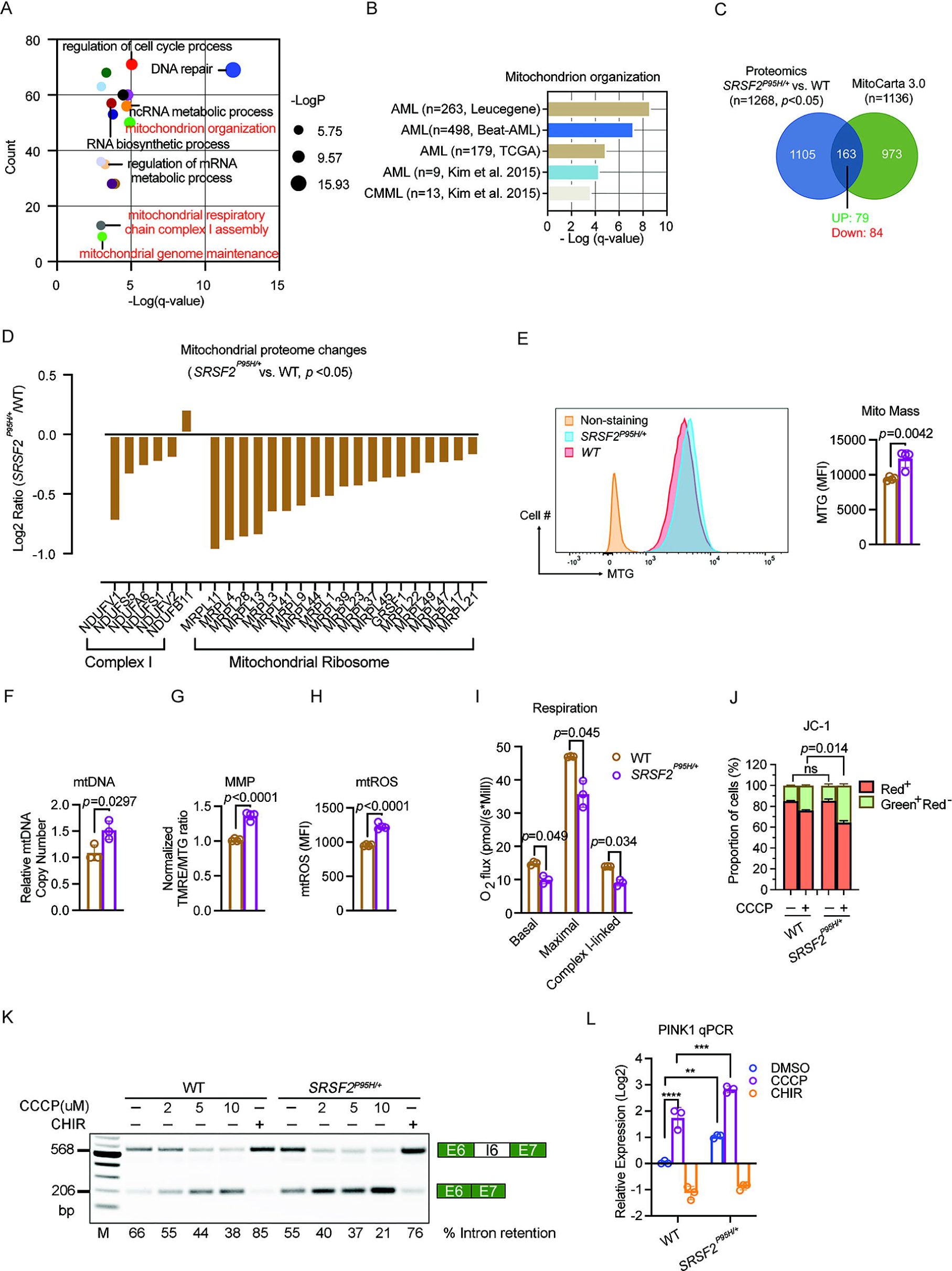
The *SRSF2* mutation is associated with accumulation of defective mitochondria. (**A**) Bubble plot for gene ontology enrichment analysis of differentially spliced genes in *SRSF2* mutant cells versus WT K562 cells. The y-axis shows the number of genes, whereas the x-axis denotes -log(q-value). The size of each circle represents the -logP value. (**B**) Bar graph for gene ontology enrichment analysis of differentially spliced genes in *SRSF2* mutant cells versus *SRSF2^+/+^*primary CMML and AML patient samples shows significantly dysregulated pathway that is associated with mitochondrion organization, with Fisher’s exact test -log[q-value] on X-axis. (**C**) Venn diagram showing overlap of proteins differentially expressed in *SRSF2^P95H/+^* versus *WT* K562 cells and MitoCarta 3.0 database. (**D**) Bar graph shows Log2 ratio of selected mitochondrial protein in *SRSF2^P95H/+^* vs. *SRSF2^+/+^* K562 cells. (**E**-**H**) Quantification of mitochondrial parameters in *WT* and *SRSF2^P95H/+^* cells (mean ± SD). **E**, MTG. **F**, Mitochondrial DNA copy number. Data are shown as ratio of mitochondrial DNA (mt-ND1) to nuclear DNA (B2M) in *wt* and *SRSF2^P95H/+^* cells. **G**, MMP per mitochondrion. **H**, mitochondrial reactive oxygen species (mtROS). (**I**) Basal, maximal, and complex I-linked mitochondrial respiratory capacity was measured in *WT* and *SRSF2^P95H/+^* cells. Maximal respiratory capacity was measured after FCCP injection. Complex I-linked respiration was measured by sequential addition of the complex I-linked substrates pyruvate, malate, and glutamate (P/M/G) and ADP. Oxygen flux expressed as respiration per million cells [pmol/(s·10^6^ cells], mean ± SD of N = 3 independent cultures. Each sample was measured in duplicate. *p* values were determined by Student’s *t* test (**E**-**I**). (**J**) Analysis of mitochondrial depolarization in *WT* and *SRSF2^P95H/+^* cells treated with DMSO or 2µM CCCP for 24h using JC-1 staining, in which a high Red^+^:Green^+^ ratio indicates high MMP (n=3). Statistical analysis was performed using a two-tailed Chi-squared test. (**K**) Representive RT-PCR results of *PINK1* splicing in *WT* and *SRSF2^P95H/+^* cells treated with DMSO, 3µM CHIR or indicated concentrations of CCCP for 24h. (**L**) RT-qPCR analysis of *PINK1* mRNA levels in *WT* and *SRSF2^P95H/+^* cells treated with DMSO, 3µM CHIR, or 2µM CCCP for 24 h (mean ± SD). \*\**p* < 0.01, \*\*\**p* < 0.001, and \*\*\*\**p* < 0.0001 (2-way ANOVA with Sidak’s multiple comparisons test).

To further assess the consequences of *SRSF2* mutation on the cellular proteome in myeloid neoplasms, we performed quantitative mass spectrometry (qMS) with a focus on the mitochondrial proteome by using human proteome (UniprotKB) and mitochondrial (MitoCarta 3.0) databases for protein identification (Figure 6C). We identified 163 mitochondrial proteins whose levels are affected by *SRSF2* mutation (Supplemental Table 5). The 84 proteins that showed significantly reduced levels in *SRSF2^P95H/+^* cells included multiple mitochondrial ribosomal proteins (Figure 6D). Of the 6 respiratory mitochondrial complex 1 proteins detected by qMS, 5 were significantly downregulated by *SRSF2* mutation (Figure 6D).

Evaluation of the mitochondrial properties of *SRSF2^P95H/+^* cells revealed elevated mitochondrial mass compared to *SRSF2^+/+^* cells based on MTG staining (Figure 6E) and immunostaining of the outer membrane protein TOMM20 (Supplemental Figure 6C). In contrast, mitochondrial mass was not increased in *SF3B1^K700E/+^* cells (Supplemental Figure 6D). Overexpression of *SRSF2^P95H^* also increased mitochondrial content in K562, TF-1, and primary AML cells when compared with the overexpression of *WT SRSF2* (Supplemental Figure 6E-G). Mitochondrial DNA (mtDNA) copy number, which correlates with mitochondrial biogenesis, significantly increased (Figure 6F), suggesting that elevated mitophagy in *SRSF2^P95H^* cells is balanced by increased mitochondrial biogenesis. This dynamic ultimately leads to increased mitochondrial content. Mitochondrial membrane potential (MMP) per unit mitochondrial mass as assessed by tetramethylrhodamine ethyl ester (TMRE)/MTG ratio and mitochondrial reactive oxygen species (mtROS) levels were significantly higher in *SRSF2^P95H/+^* cells (Figure 6G and 6H). This chronic oxidative stress may damage the mitochondria and mtDNA. To test mitochondrial function, we subjected parental and *SRSF2^P95H/+^* cells to high resolution respirometry. Surprisingly, both the basal and maximal respiratory capacity were significantly lower in *SRSF2^P95H/+^* cells (Figure 6I). As *SRSF2* mutation dramatically altered the splicing of complex I related genes and reduced complex I protein levels, we further tested complex I respiratory capacity. Complex I-linked respiration, which was measured by sequential addition of the complex I-linked substrates pyruvate, malate, and glutamate (P/M/G) and ADP, was also significantly lower in *SRSF2^P95H/+^* cells relative to *SRSF2^+/+^* cells (Figure 6I). The elevated mitochondrial mass and MMP in *SRSF2^P95H/+^* cells therefore could not maintain high respiratory capacity. These functional data together with the increase in mitophagy demonstrate a marked defect in mitochondrial function associated with the leukemogenic *SRSF2^P95H^* mutation.

We next treated *WT* and *SRSF2^P95H/+^* K562 cells with a mitochondrial uncoupler, carbonyl cyanide m-chlorophenylhydrazone (CCCP) and then measured mitochondrial depolarization occurring during apoptosis using JC-1 staining (33). Although *WT* cells responded to CCCP treatment with reduced MMP, the *SRSF2* mutation further decreased MMP, suggesting a higher demand for efficient mitophagy in response to mitochondrial stress in *SRSF2^P95H/+^*cells (Figure 6J). Overall, these data support a model in which splicing defects associated with the *SRSF2* mutation disrupt mitochondrial function and lead to compensatory increased mitochondrial content and turnover.

As *SRSF2^P95H^* increases the abundance of *PINK1* mRNA, we further hypothesize that mitochondrial dysfunction signals to the splicing apparatus to promote excision of intron-6, stabilizing *PINK1* mRNA to meet an increased demand for mitophagy. In support of this hypothesis, addition of CCCP promotes excision of intron-6 in a dose-dependent manner in cells with wild-type *SRSF2* and to an even greater extent in *SRSF2^P95H/+^*cells (Figure 6K). Consistent with the destabilizing impact of the PTC in intron-6, this shift in splicing was associated with an increase in *PINK1* mRNA abundance (Figure 6L). This response in cells with wild-type *SRSF2* suggests a general mechanism, independent of the *SRSF2* mutation, for sensing mitochondrial stress through modulation of *PINK1* splicing, leading to increased expression of *PINK1* mRNA and protein.

### Increased mitophagy is a therapeutic vulnerability in *SRSF2* mutant hematologic malignancies

GSK-3i reduces *PINK1* expression, impairs mitophagy, and is selectively lethal to *SRSF2^P95H/+^* cells. Given that PINK1-mediated mitophagy is required for the survival of normal hematopoietic stem cells (HSCs) (34) and acute myeloid leukemia stem cells (LSCs) (35, 36), we hypothesized that a greater dependency on PINK1-mediated mitophagy upon stress may be the basis for the selective toxicity of GSK-3i to *SRSF2^P95H/+^* cells. To explore this hypothesis, we evaluated the effects of GSK-3i on mitochondria. GSK-3i increased MMP (Figure 7A) and promoted accumulation of mitochondrial mass in K562 cells with *WT SRSF2* (Figure 7B and Supplemental Figure 7A), consistent with previous reports in mouse cardiomyocytes and HSCs (37, 38). GSK-3i in *SRSF2^P95H/+^*cells impaired mitophagy (Figure 5B and 5C) and further increased MMP (Figure 7A) and mitochondrial content (Figure 7B and Supplemental Figure 7A-D) compared to *WT* counterparts. Gene set enrichment analysis (GSEA) revealed that GSK-3i with CHIR strongly enriches for genes representing enhanced mitochondrial biogenesis and oxidative phosphorylation (OXPHOS), a mitochondrial stress that may increase the requirement for PINK1-mediated mitochondrial surveillance (Figure 7C).

**Figure 7.**
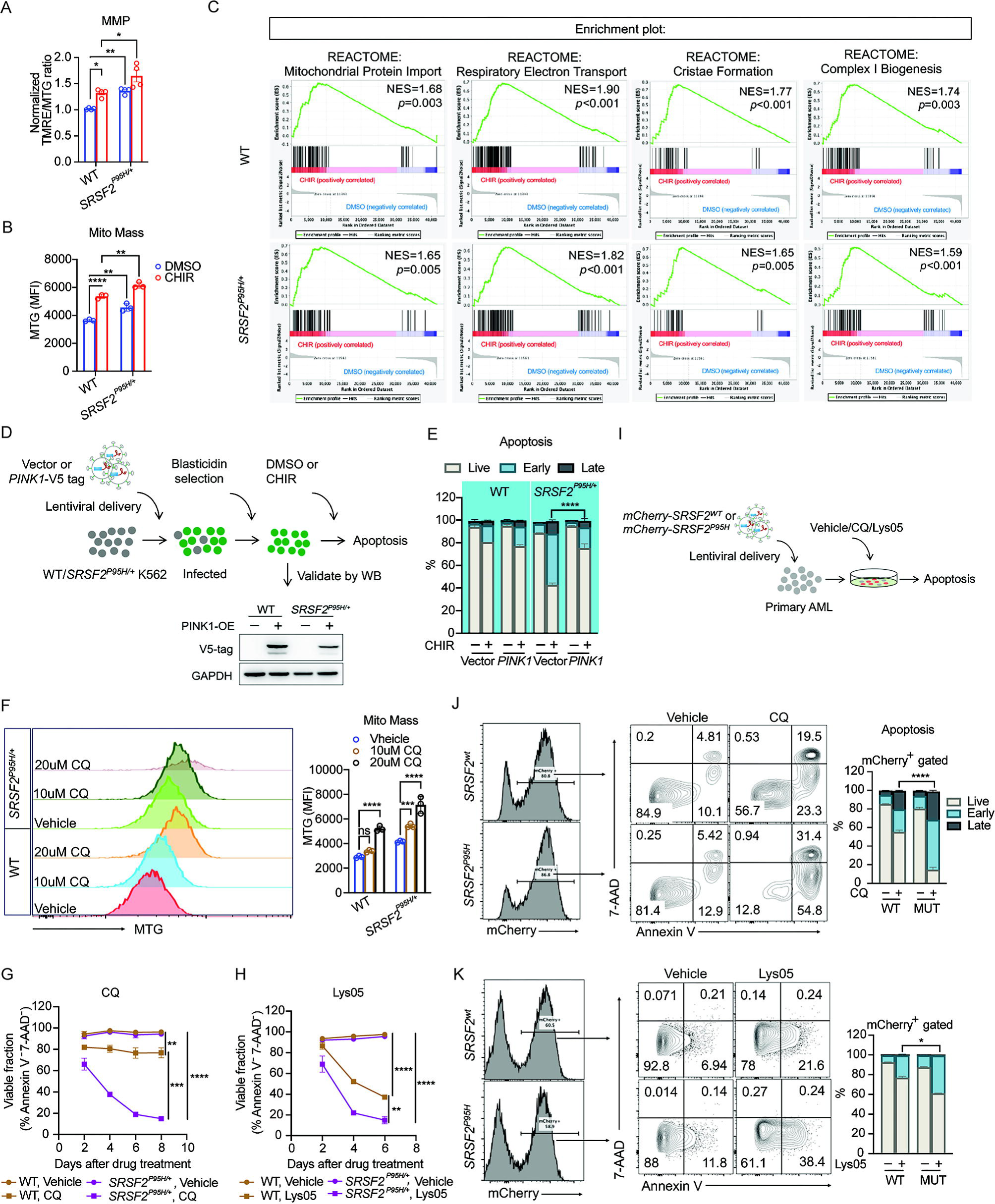
Targeting mitophagy in *SRSF2* mutant hematologic malignancies. (**A**-**B**) Quantification of mitochondrial parameters in *WT* and *SRSF2^P95H/+^*cells treated with DMSO or 3µM CHIR. **A**, MMP per mitochondrion. **B**, MTG. (**C**) GSEA showing mitochondrial related enrichment plots for CHIR versus DMSO treated *WT* and *SRSF2^P95H/+^* cells. (**D**) Schematic of generating *WT* and *SRSF2^P95H/+^* stable lines overexpressing *PINK1* for apoptosis assay with or without CHIR. (**E**) Percentages of viable, early-, and late-apoptotic cells in *PINK1* overexpressing *WT* and *SRSF2^P95H/+^*cells treated with DMSO or 3µM CHIR *in vitro* for 8d. (**F**) Quantification of mitochondrial mass by MTG staining in *WT* and *SRSF2^P95H/+^* cells treated with vehicle or indicated concentrations of CQ for 6d. (**G**-**H**) Percentage of viable cells based on 7-AAD and Annexin V flow cytometric analysis of *WT* and *SRSF2^P95H/+^* cells treated with vehicle, 20µM CQ (**G**), or 2uM Lys05 (**H**) *in vitro*. (**I**) Schematic representation of lentiviral delivery of *SRSF2^wt^* or *SRSF2^P95H^* to primary AML cells for Chloroquine and Lys05 treatment followed by apoptosis assay. (**J**-**K**) Representative flow cytometric analysis (left) and quantification (right) of apoptosis in primary cells from patient with AML overexpressing either *WT* or *SRSF2^P95H^*, as measured by Annexin V and 7-AAD staining in absence or presence of 15µM Chloroquine (**J**) or 2uM Lys05 (**K**) for 4d. Data in A, B, F, G and H are presented as the mean ± SD. \**p* < 0.05, ** *p* < 0.01, *** *p* < 0.001, and **** *p* < 0.0001 (2-way ANOVA with Sidak’s multiple comparisons test). For data in E, J, and K, statistical analysis was performed using a two-tailed Chi-squared test.

To test directly whether GSK-3i-induced alternative splicing of *PINK1* causes cell death in *SRSF2^P95H/+^* cells, we expressed a *PINK1* cDNA lacking intron-6 and treated with CHIR (Figure 7D). Expression of full length *PINK1* completely rescued the cell death caused by GSK-3i in *SRSF2^P95H/+^* cells (Figure 7E), demonstrating an essential role for *PINK1* mis-splicing in the preferential cytotoxicity of GSK-3i in *SRSF2* mutant cells.

Our data indicate that increased mitophagy is a targetable vulnerability in *SRSF2^P95H/+^* cells. We tested this hypothesis further by inhibiting mitophagy downstream of PINK1 by treating *SRSF2^+/+^* and *SRSF2^P95H/+^* cells with the lysosomal inhibitor CQ. Inhibiting mitophagy with CQ caused accumulation of mitochondria (Figure 7F) and preferentially killed *SRSF2^P95H/+^* cells (IC_50_ = 14.1µM) compared to *SRSF2^+/+^* cells (IC_50_ = 29.1µM) (Figure 7G, and Supplemental Figure 7E). Preferential sensitivity of *SRSF2^P95H/+^* cells to autophagy inhibitors was further validated with Lys05 (39, 40), a potent dimeric form of CQ (Figure 7H and Supplemental Figure 7F). Next, we introduced *SRSF2^P95H^*or *SRSF2^wt^* into *SRSF2^+/+^* primary AML cells along with the mCherry reporter (Figure 7I and Supplemental Table 1). Expression of *SRSF2^P95H^* resulted in increased sensitivity to CQ (Figure 7J) and Lys05 (Figure 7K) compared to primary AML cells expressing *SRSF2^wt^*.

Together, these data support the hypothesis that aberrant splicing of *PINK1* caused by GSK-3i reduces *PINK1* expression and inhibits clearance of defective mitochondria in *SRSF2* mutant cells, disrupting the homeostatic balance required for survival. Targeting mitophagy therefore represents a promising therapeutic strategy to eradicate *SRSF2* mutant hematologic malignancies (Figure 8).

**Figure 8.**
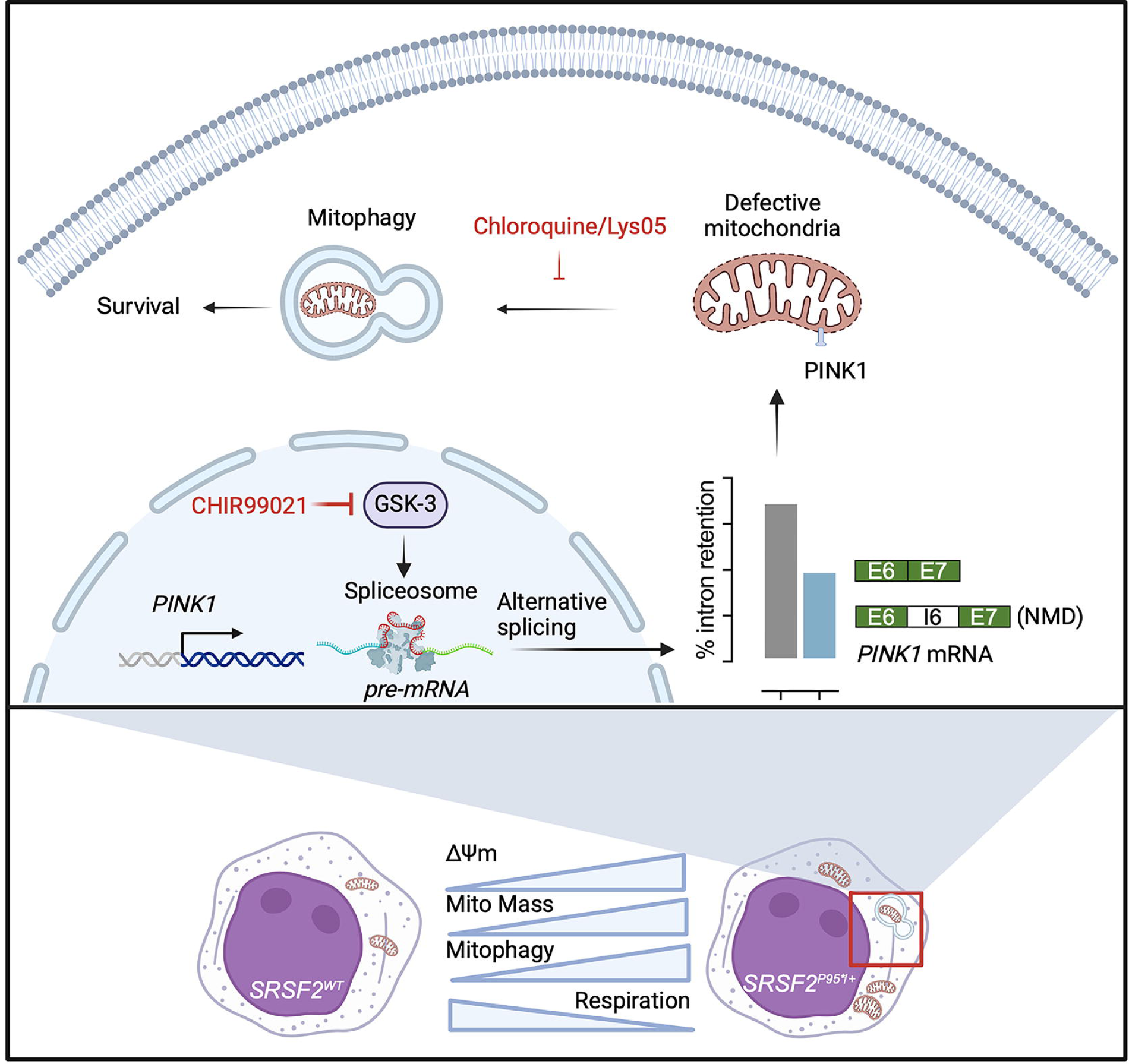
Proposed model for dependency of *SRSF2* mutant leukemias on PINK1-mediated mitophagy and preferential cytotoxicity of GSK-3i and mitophagy inhibitor Chloroquine and Lys05.

## DISCUSSION

Splicing factor mutations are common in MDS and AML (2, 41, 42), but how these mutations alter cellular function remains unclear. The results presented here show a marked increase in mitophagy specifically in *SRSF2* mutant MDS and AML compared to other forms of AML and MDS. The recurrent *SRSF2^P95H^* mutation alters the splicing of mRNAs encoding mitochondrial proteins, disrupts mitochondrial function, and increases mitochondrial mass and turnover, reflecting a dependency on a higher level of mitophagy for survival. Our results identify a mechanism of mitochondrial surveillance in which mitochondrial dysfunction enhances the splicing of *PINK1* mRNA to a more stable form, leading to increased abundance of *PINK1* mRNA and protein. Inhibition of GSK-3 impairs *PINK1* splicing, reduces mitophagy, and selectively kills cells with the *SRSF2^P95H^*mutation. Inhibition of mitophagy by targeting lysosomal function is also lethal to *SRSF2^P95H/+^* cells. These data therefore reveal a dependency of splicing factor mutant MDS and AML and identify potential therapeutic targets for these hematologic malignancies.

The altered splicing of nuclear encoded mitochondrial mRNAs observed here fits well with previous work identifying altered splicing of mRNAs involved in mitochondrial function in primary cells from MDS and AML patients with the *SRSF2^P95^* mutation (7, 9, 32). We also find that markers of mitophagy are robustly increased in AML and MDS patients with the *SRSF2^P95/+^* mutation in the TCGA dataset. Mitochondrial turnover is required for self-renewal of normal HSCs (34, 43, 44) and leukemic stem cells (36, 44, 45), and was recently shown to mediate resistance to BH3 mimetics in AML cells (46–49). Thus, while inhibition of mitophagy or autophagy has been shown to be toxic to AML cells (36, 40, 45, 46), a mechanism for increased mitophagy in AML has not previously been defined. Our data demonstrate significantly greater sensitivity to mitophagy/autophagy inhibitors in *SRSF2^P95/+^* cells, reveal a mechanism connecting this leukemogenic splicing factor mutation to mitochondrial dysfunction and increased mitophagy, and identify *PINK1* splicing as a targetable vulnerability in *SRSF2* mutated AML and MDS.

The data presented here reveal a homeostatic mechanism for the regulation of mitochondrial clearance. Prior work has established that, under basal conditions, PINK1 protein is degraded within mitochondria with a high MMP, whereas low MMP associated with mitochondrial dysfunction allows PINK1 protein stabilization and initiation of mitophagy (50). The homeostatic mechanism described here functions by altering *PINK1* splicing: mitochondrial dysfunction, whether caused by direct pharmacological disruption of the MMP or indirectly by the *SRSF2* mutation, promotes the excision of a poison intron to yield a more stable *PINK1* mRNA. Although the *SRSF2^P95H^*mutation could directly alter the splicing of *PINK1* mRNA, we propose that the mechanism is indirect. SRSF2 typically regulates cassette exon selection whereas we observe enhanced intron excision and, importantly, CCCP alters *PINK1* splicing in cells with wild-type *SRSF2,* supporting a homeostatic mechanism that is sensitive to but independent of the *SRSF2* mutation. Although the altered splicing of *PINK1* mRNA described here is distinct from the well-established regulation of PINK1 protein stability by MMP, both mechanisms result in increased PINK1 protein abundance under conditions of mitochondrial stress. Our data thus support a mechanism for sensing mitochondrial stress through modulation of *PINK1* splicing, increasing *PINK1* expression to support an increased demand for mitophagy in the setting of mitochondrial dysfunction.

Splicing out of the poison intron requires GSK-3, which phosphorylates multiple core splicing factors, including SRSF2, and regulates splicing at a transcriptome-wide level in diverse cell types (22–26). Thus, pharmacologic inhibition or *GSK3* knock out impairs splicing of *PINK1*, leading to retention of the poison intron and reduction in overall levels of *PINK1* mRNA and protein. Altered splicing of *PINK1* explains the lethality of GSK-3 inhibitors in *SRSF2^P95H/+^* cells, as survival in the presence of a GSK-3 inhibitor is rescued by expression of *PINK1* cDNA. Furthermore, chloroquine and Lys05, which target autophagy downstream of *PINK1,* also preferentially kill *SRSF2^P95H/+^* cells, supporting the hypothesis that these splicing factor mutant cells are dependent on mitophagy for survival and suggesting an actionable therapeutic target in splicing factor mutant myeloid neoplasms.

Currently there are no FDA approved drugs targeting splicing factor mutant malignancies. Here we identify mitophagy as a therapeutic vulnerability specifically in AML and MDS driven by hotspot mutations in the splicing factor *SRSF2*. We have also uncovered a mechanism for mitochondrial surveillance that is mediated through GSK-3-dependent alternative splicing of *PINK1* and show significantly increased sensitivity to GSK-3 or autophagy inhibition in splicing factor mutant cells. Several GSK-3 inhibitors are either in wide clinical use or have been shown to be safe in phase I-III clinical trials (51–54) and could be repurposed to treat MDS. Hence targeting mitophagy with GSK-3 inhibitors or general autophagy inhibitors may provide a new opportunity to treat *SRSF2* mutant MDS and AML.

## MATERIALS AND METHODS

### Sex as a biological variable

This study used de-idenitified primary patient cells from male and female patients. However, sex was not considered as a biological variable because the study was not powered to distinguish sex differences. Xenografts of human cells into murine hosts were performed using female mice only. Sex of the host was not considered as a biological variable in these xenograft experiments.

### Cell lines, primary human samples, constructs and nucleofection

BM- or peripheral blood-derived MNCs from AML or CMML patients were obtained from the Stem Cell and Xenograft Core Facility at University of Pennsylvania (RRID: SCR_010035). Detailed patient characteristics are listed in Supplemental Table 1. CD34^+^ cells were purified from CMML patients through immunomagnetic selection (Miltenyi Biotec) according to the manufacturer’s instructions. CD34^+^ cells from healthy donors were purchased from STEMCELL Technologies. Cryopreserved AML MNCs were resuspended in IMDM medium supplemented with 15% BIT (bovine serum albumin, insulin, transferrin, STEMCELL Technologies), 100ng/ml SCF, 50ng/ml FLT3L, 20ng/ml IL-3, and 20ng/ml G-CSF. CD34^+^ cells enriched from CMML patients or healthy donors were cultured in StemSpan SFEM II medium (STEMCELL Technologies) supplemented with 10% fetal bovine serum (FBS), 1% L-glutamine, 10ng/ml IL-3, 10ng/ml IL-6, and 25ng/ml SCF. TF-1 cells were maintained in RPMI supplemented with 10% FBS and 2ng/ml hGC-SCF (PeproTech). K562 cells were cultured in IMDM supplemented with 10% FBS. Cells were maintained at 37°C and 5% CO_2_. *PINK1* (pLenti6-DEST PINK1-V5 WT, 13320), *shUPF1* (PLKO.1-UPF1-CDS, 136037), *WT* (pRRL_SRSF2_WT_mCherry, 84020), and *P95H SRSF2* (pRRL_SRSF2_P95H_mCherry, 84023) lentiviral overexpression constructs were purchased from Addgene. Lentiviruses were produced and used to transduce TF-1, K562, and primary AML cells, as described previously (55). 2X10^6^ primary AML cells were transduced with WT and *P95H SRSF2* lentivirus by spin infection in growth medium containing 4% LentiBlast (OZ Biosciences). Transduction efficiency was 30-60%. HEK293T cells were cultured in DMEM supplemented with 10% FBS, and 1% penicillin/streptomycin. *GSK3A/B* DKO HEK293T cells were generated by lentiviral delivery of Cas9 and guide RNA sequence targeting *GSK-3A* (GCCTAGAGTGGCTACGACTG) and *GSK-3B* (AGATGAGGTCTATCTTAATC) followed by single clone selection as previously described (51). *GSK3A/B* DKO HEK293T cells were transfected using Lipofectamine 3000 (Invitrogen) according to manufacturer’s instructions.

### Flow cytometry, apoptosis assay and cell cycle analysis

For flow cytometric apoptosis assay, K562, TF-1, primary AML blast or CD34^+^ cells purified from CMML patients or healthy donors were suspended in Annexin V Binding buffer (BioLegend #422201) and incubated with anti-Annexin V antibody (BioLegend #640920) and 7-AAD (BioLegend #420403) for 15 min at room temperature (RT) in the dark. Immunophenotypes of viable cells or cells in early apoptosis, or late apoptosis were defined as Annexin V^-^7-AAD^-^, Annexin V^+^7-AAD^-^ or Annexin V^+^7-AAD^+^, respectively. For cell cycle analysis, cells were fixed and permeablized in 70% ethanol in -20°C for 1 h, washed in cold FACS buffer, and then stained with anti-Ki67 (BD #350506) for 30 mins on ice. After staining, cells were washed twice in FACS buffer and resuspended in FACS buffer containing 10µM DAPI (BioLegend #422801). Stained cells were then tested by flow cytometry. Differences in apoptosis between culture conditions were analyzed by two-sided chi-squared test.

### Cell growth assay

Triplicates of WT and *SRSF2^P95H/+^* K562 cells were seeded at 10,000 cells per well in 96-well plates and treated with indicated concentrations of CHIR (Cayman #13122) or Chloroquine (Sigma #C6628). Four days after culture, MTS reagent (Abcam #ab197010) was added to cell medium at a final concentration of 0.5 mg/mL for 3 h at 37°C, and cell viability was measured as per manufacturer’s instructions. IC_50_ was determined with GraphPad Prism 8 using baseline correction (by normalizing to vehicle control), the asymmetric (five parameter) equation and least squares fit.

### Immunoblotting

Cells were lysed in RIPA buffer plus protease inhibitor cocktail (Sigma, P8340), phosphatase inhibitor cocktail #2 (Sigma #P5726) and #3 (Sigma #P0044) used 1:100 each. Supernatants were collected after centrifugation at 14,000 rpm for 15 min at 4°C, adjusted to 1x laemmli sample buffer and subjected to SDS-PAGE and then immunoblotted as described previously (51). Antibodies for biochemical studies purchased from Cell Signaling Technology included: anti-GAPDH (#2118), cleaved caspase 8 (#9496), cleaved caspase 3 (#9661), Flag-tag (#14793), V5-tag (#13202), and β-catenin (#9562). Other antibodies included antibodies to PINK1 (Invitrogen #PA5-86941) and β-actin (Sigma #A5441).

### Real time qPCR and RT-PCR

RNA was extracted from TF-1, K562, and primary cells using Rneasy Kit (Qiagen) according to the manufacturer’s instructions. cDNA was synthesized using the SuperScript III (Life Technologies) according to the manufacturer’s instructions. For detection of PINK1 mature mRNA, the following primers were used: forward: 5’-GCCTCATCGAGGAAAAACAGG-3’; reverse: 5’-GTCTCGTGTCCAACGGGTC-3’. For detection of PINK1 pre-mRNA, the following primers were used: intron 5 forward: 5’-CCTTTGCCTGGGGATTTTGC-3’; intron 5 reverse: 5’-GGGGCTAATGGCTCAGTGTT-3’; Intron 1 forward: 5’-GAGGCGAGGGTCCTTAAAGC-3’; intron 1 reverse: 5’-TGCGACAGGAGCTGTAATCG-3’. For detection of the PINK1 aberrant junction, the following primers were used: forward: 5’-GGTGATCGCAGATTTTGGCT-3’; reverse: 5’-GCCCGAAGATTTCATAGGCG-3’. The primers used for *BCL-X* isoform specific real-time qPCR analysis: *BCL-X* all forward: 5’-CATCAATGGCAACCCATCCTG-3’; reverse: 5’-GCAGTTCAAACTCGTCGCCT-3’; *BCL-Xs* forward: 5’-GAGCTTTGAACAGGATACTTTTGTGG-3’: reverse: 5’-TTCCGACTGAAGAGTGAGCC-3’. The primers used for *DEPDC1* isoform specific qPCR analysis: *DEPDC1* all forward: 5’-TGATGCAATGGGTACGAGGT-3’; reverse: 5’-TCTTCCAGCAAGAAGCTCATCA-3’; *DEPDC1* long isoform forward: 5’-GAACTCGGAGAGTCTAGTGCC-3’; reverse: 5’-CATCGATGGCAACCCTCTCT -3’. The efficiency of *UPF1* knockdown was measured at the mRNA level by RT-qPCR: forward, 5’-AATTTGGTTAAGAGACATGCGG -3’; reverse, 5’-TCAGGGACCTTGATGACGTG-3’.

### Mitophagy measurement and fluorescence microscopy

*WT* and *SRSF2^P95H/+^* K562 cells cells were treated with DMSO or 3µM CHIR for 44 h, then seeded on poly-D-lysine coated slides for 4 h at 37°C. After incubation, cells were fixed with 4% paraformaldehyde (pH 7.4) for 15 mins, permeabilized with 0.1% Triton X-100 for 15 mins and blocked with 2% bovine serum albumin in PBS for 1 h at room temperature. Cells were immunostained with mouse anti-TOMM20 antibody (1:100 dilution, clone 4F3, Sigma-Aldrich #WH0009804M1), rabbit anti-LAMP1 antibody (1:100 dilution, ABclonal #A2582) overnight at 4°C in a humidified chamber. Secondary antibodies in anti-rabbit AF568 (Invitrogen #A11011) and anti-mouse AF488 (Invitrogen #A11001, 1:1000 dilution) were incubated at room temperature for 60 mins. The slides were then air-dried in dark at room temperature for 10 minutes and mounted in anti-fade mounting medium (Abcam #ab104135) before confocal imaging. Co-localization of TOMM20 and LAMP1 was quantified with JaCoP plug-in in Fiji. Flux analysis was performed to quantify the turnover of mitochondria in lysosomes by evaluating the accumulation of mitochondria in the presence of the lysosomal inhibitors (31). Cells were cultured in the presence or absence of 100uM Chloroquine or 50uM Lys05 for 4 hours following which cells were subjected to mito-tracker green (MTG) (Invitrogen #M7514) staining for flow cytometric analysis. Mitophagic flux was determined by subtracting the MTG value for untreated cells from the value for cells treated with either CQ or Lys05 (31, 40).

For PINK1 staining, cells were immuno-stained with rabbit anti-PINK1 antibody (1:100 dilution, Proteintech #23274-1-AP), mouse anti-TOMM20 antibody (1:200 dilution), and mouse anti-PARKIN antibody (1:100 dilution, Proteintech #66674-1-1g) as previously described. For high resolution imaging, the stained slides were imaged using Zeiss LSM880 confocal microscope with Airyscan. Images were captured using a 60X objective with Z stacks followed by deconvolution analysis.

### Measurements of mitochondrial mass, membrane potential, ROS and mitochondrial DNA copy number

For mitochondrial mass measurement, cells were stained with 15nM MTG for 20 mins at room temperature in dark followed by flow cytometric analysis. For mitochondrial membrane potential analysis, cells were stained with 200nM TMRE (Invitrogen #T669) for 20 mins at room temperature. JC-1 (Invitrogen #M34152) staining was used for detecting mitochondrial depolarization occurring in apoptosis. WT and *SRSF2^P95H/+^*K562 cells were treated with or without 2µM CCCP for 24h, and then stained with 1µM JC-1 for 20 mins. Cellular ROS levels were measured by FACS following cell surface staining using the CellROX kit (Invitrogen, C10444) according to manufacturer’s instructions. To quantify mitochondrial DNA copy number, quantitative PCR was performed with primers for mitochondrial mt-ND1 (Forward: GCAGAGACCAACCGAACCCCC; Reverse: GGGCCTGCGGCGTATTCGAT) and nuclear β2 microglobulin (B2M) (Forward: GCAGAGACCAACCGAACCCCC; Reverse: GGCGGGCCACCAAGGAGAAC).

### Transmission electron microscopy

K562 WT and *SRSF2^P95H/+^* cells for electron microscopic examination were fixed with 2.5% glutaraldehyde, 2.0% paraformaldehyde in 0.1M sodium cacodylate buffer, pH7.4, overnight at 4°C. After subsequent buffer washes, the samples were post-fixed in 2.0% osmium tetroxide for 1 hour at room temperature and rinsed in dH_2_O prior to en bloc staining with 2% uranyl acetate. After dehydration through a graded ethanol series, the tissue was infiltrated and embedded in EMbed-812 (Electron Microscopy Sciences, Fort Washington, PA). Thin sections were stained with uranyl acetate and lead citrate and examined with a JEOL1010 electron microscope fitted with a Hamamatsu digital camera and AMT Advantage image capture software. Images of single cells were saved as separate image files and an observer blind to the identity of each cell counted vesicles per cell.

### Measurement of mitochondrial respiration

Oxygen consumption was measured using high-resolution respirometry Oxygraph-2k (Oroboros Instruments, Innsbruck, Austria) with a polarographic oxygen electrode and two 2-mL chambers allowing for parallel measurements^52^. Briefly, K562 WT and *SRSF2^P95H/+^* cells in MiR05 buffer (0.5mM EGTA, 3mM MgCl2, 60mM lactobionic acid, 20mM taurine, 10mM KH2PO4, 20mM HEPES, 110 mM D-sucrose, 1G/L BSA) were added into the closed chamber through a small capillary tube. Oxygen concentration (μmol/L) and oxygen flux [pmol/(s×Mill)] were simultaneously recorded in real-time. During the assay, 5ug/ml Digitonin, 5mM Pyruvate, 2mM Malate, 10mM Glutamate, 2.5mM ADP, 5nM Oligomycin, 0.5uM FCCP, 0.5uM Rotenone, and 2.5uM Antimycin A were added sequentially.

### K562 xenograft model

5X10^6^ K562 WT or *SRSF2^P95H/+^* cells were subcutaneously implanted into the flank of female NSG mice of 6–8 weeks of age. Mice were then treated with 30mg/kg body weight CHIR or vehicle (10% DMSO, 45% PEG400, and 45% PBS, injected subcutaneously) daily for the duration of the implantation period. Tumor size was measured three times a week using a caliper. Tumor volume was calculated by using the ellipsoid formula: (length X width2)/2.

### RNA-seq sample preparation

K562 isogenic lines were treated with 3µM CHIR for 24 h. After treatment, the cells were washed with cold PBS, and RNA was isolated using Rneasy Kit (Qiagen) according to the manufacturer’s instructions. cDNA library preparation, sequencing, and raw read filtering methods were described previously (22).

### Quantification of RNA-seq data

Raw sequence reads (10^8^ reads per replicate sample) were aligned to the human reference sequence hg38 by STAR 2.4.2a using two-pass alignment. Raw gene counts were compiled into total gene counts, then analyzed using EdgeR 3.20.2 to assess the significance of changes between cohorts. Gene changes with a padj value < 0.05 were considered significant. Significant gene expression changes are provided in Supplemental Table 2. Heatmaps for BCL2 family gene expression and mitophagy related genes expression were generated by using Morpheus tool (Broad Institute, https://software.broadinstitute.org/morpheus/). Briefly, normalized expression values were transformed to z-scored log2 expression by subtracting row mean and then dividing by row standard deviation. RNA-seq data were deposited in GEO (GSE235600) and are accessible as described below.

### Identification of differential splicing events

Alternative splicing analyses relied on RNA-Seq reads mapped to the reference human genome as described using rMATS v4.1.1 with the default parameters (28). Events were defined as significant if (1) the FDR corrected p value was smaller than 0.05 and (2) dPSI was larger than 10%. Examples of splicing and mis-splicing events were visualized with IGV (Broad Institute). Heatmap for alternative splicing events between *WT* or *SRSF2^P95H/+^* cells was generated by using Morpheus. Hierarchical clustering was performed using metric one minus person correlation.

### Gene ontology analysis

Gene ontology analysis was performed using Metascape (https://metascape.org) using the KEGG, and GO specific signatures according to the manual.

### GSEA analysis

GSEA analysis was performed using GSEA version 4.3 (Broad). Normalized reads values produced from gene expression analysis were formatted into GCT files containing expression values for genes in different biological states. CLS files were manually built to label biological states involved in each study. Following parameters were used: Number of permutations = 1000, permutation type = gene_set, chip platform = human_Ensembl_gene_ID_MSigDB.v2023.1.chip. Other parameters were used at default values.

### Mass spectrometry

*SRSF2^+/+^* (*WT)* and *SRSF2^P95H/+^* K562 cell pellets were resuspended in a solution of 100 mM ammonium bicarbonate and 8 M urea. Samples were sonicated for 15 sec then placed on ice for 15 sec ten times, and then were centrifuged at 20,000xg for 5 min at 4°C. Protein concentration in supernatants was measured by BCA assay. For each sample, 100 µg of protein was diluted to 50 µL at a final concentration of 10 mM DTT and incubated at 56°C for 30 min to reduce cystines to cysteines. Alkylation was performed by adding 5.5 µL of 0.5 M iodoacetamide and incubating at RT in the dark for 40 minutes. Samples were diluted to 250 µL with 50 mM Tris-HCl (pH 8.3), 2 µg of sequencing grade modified trypsin (Promega) was added, and samples were incubated at 37°C overnight. Samples were acidified to 0.1% trifluoroacetic acid (TFA) and immediately desalted using C18 stagetips, washing with 0.1% formic acid and eluting with 0.1% formic acid in 60% acetonitrile (CAN). Peptides were dried in a Savant SpeedVac and then resuspended in 20 uL of 0.1% trifluoroacetic acid (TFA). UV absorption at 280 nm (A280) was measured to normalize injection volumes and samples were run on a Dionex UltiMate 3000 nanoLC and Q Exactive HF (Thermo Scientific). Peptides were loaded on a C18 trap column (Thermo Scientific), washed with buffer A (0.1% formic acid) and then separated using an analytical column (75 µm x 15 cm) packed in-house with C18 resin (Dr. Maisch, GMBH), using an analytical gradient of 5% buffer B (0.1% formic acid in 80% ACN) to 25% over 90 min, and then 25% to 45% over 30 min. MS detection was performed using data independent acquisition with 24 *m/z* windows. Data were searched using DIA-NN with a UniProt FASTA digest library-free search and without heuristic protein inference. The mass spectrometry proteomics data have been deposited to the ProteomeXchange Consortium via the PRIDE (56) partner repository with the dataset identifier PXD043213.

### AML TCGA analysis

Gene expression analysis of *OPTN*, *ULK1* and *TOMM7* in AML patients from the TCGA project (57) was retrieved from cBioportal for Cancer Genomics (https://www.cbioportal.org) (58), and visualized using Prism (GraphPad). GEPIA (http://gepia.cancer-pku.cn) was used for overall survival (OS) analysis based on high and low expression of *OPTN* in publicly available TCGA datasets.

### Statistics

Statistical analysis was performed using Prism version 8 software. Statistical differences between two groups were determined by 2-tailed Student’s *t* test. To assess the statistical significance of differences between more than two treatments, one-way or two-way ANOVA followed by Sidak’s multiple comparisons test were used. P < 0.05 was considered significant.

### Study Approval

All mouse studies were carried out through the Stem Cell and Xenograft Core under a protocol approved by the University of Pennsylvania Institutional Animal Care and Use Committee (IACUC).

### Data availability

RNA-seq data were deposited in the NCBI Gene Expression Omnibus (59) and are accessible through GEO Series accession number GSE235600, https://www.ncbi.nlm.nih.gov/geo/query/acc.cgi?acc=GSE235600. The mass spectrometry proteomics data have been deposited to the ProteomeXchange Consortium via the PRIDE (56) partner repository with the dataset identifier PXD043213. Supporting data values for all figures are provided in the attached excel file “Supporting data values.xlsx”

### Author Contributions: (CRediT statement)

Conceptualization: XL, PSK

Methodology: XL, MQV, NS, MPC, RM, KJ

Supervision and funding acquisition: PSK, OAW, KWL, DCW

Investigation: XL, SAD, RFS, RM, KJ, MQV, OP, AAM, CL

Resources: PSK, RFS, OAW, NS, MPC, JH, KWL, DCW

Writing and editing: XL, PSK, RFS, RM, KJ, NS

Visualization: XL, OP, AAM

Supervision: PSK, OAW, KWL, DCW, MPC, JH

Funding acquisition: PSK, KWL

## Acknowledgments

We appreciate helpful suggestions from Drs. M. Celeste Simon, Brian Keith, and Vikram Paralkar. We also thank Akmal Salimov for assistance with SCXC sample characterization, and Blake Hernandez for suggestions with confocal imaging. This research was supported by grants to PSK from the National Institutes of Health (NIH, R01HL141759 and R56DK133258), the Leukemia and Lymphoma Society (#8036-23), and the Institute for Translational Medicine and Applied Therapeutics (ITMAT) at the University of Pennsylvania; to KWL from the NIH (R35-GM118048), and MPC from a Veterans Administration Merit Award (I01BX004662).

