## Supplemental data for "A mitochondrial surveillance mechanism activated by *SRSF2* mutations in hematologic malignancies"

**List of Supplemental Materials**

**Supplemental Figures 1 – 7**

**Supplemental Tables 1 – 5 (excel)**

**Supplemental Table 1** Clinical and genetic characteristics of primary AML and CMML patient samples.

**Supplemental Table 2** Differentially expressed genes in CHIR versus DMSO treated WT, *SRSF2*$^{P95H/+}$ and *SF3B1*$^{K700E/+}$ K562 cells with a cutoff of padj < 0.05 and log2[fold change]l > 1.

**Supplemental Table 3** Alternative splicing genes in CHIR versus DMSO treated WT, *SRSF2*$^{P95H/+}$ and *SF3B1*$^{K700E/+}$ K562 cells with a cutoff of false discovery rate (FDR) < 0.05 and dPSI >10%.

**Supplemental Table 4** Gene ontology (GO) analysis of differentially spliced genes in *SRSF2*$^{P95H/+}$ vs WT K562 cells.

**Supplemental Table 5** Differentially expressed mitochondrial proteins in *SRSF2*$^{P95H/+}$ versus *SRSF2*$^{+/+}$ K562 cells.

### Supplemental Figures

#### Figure S1

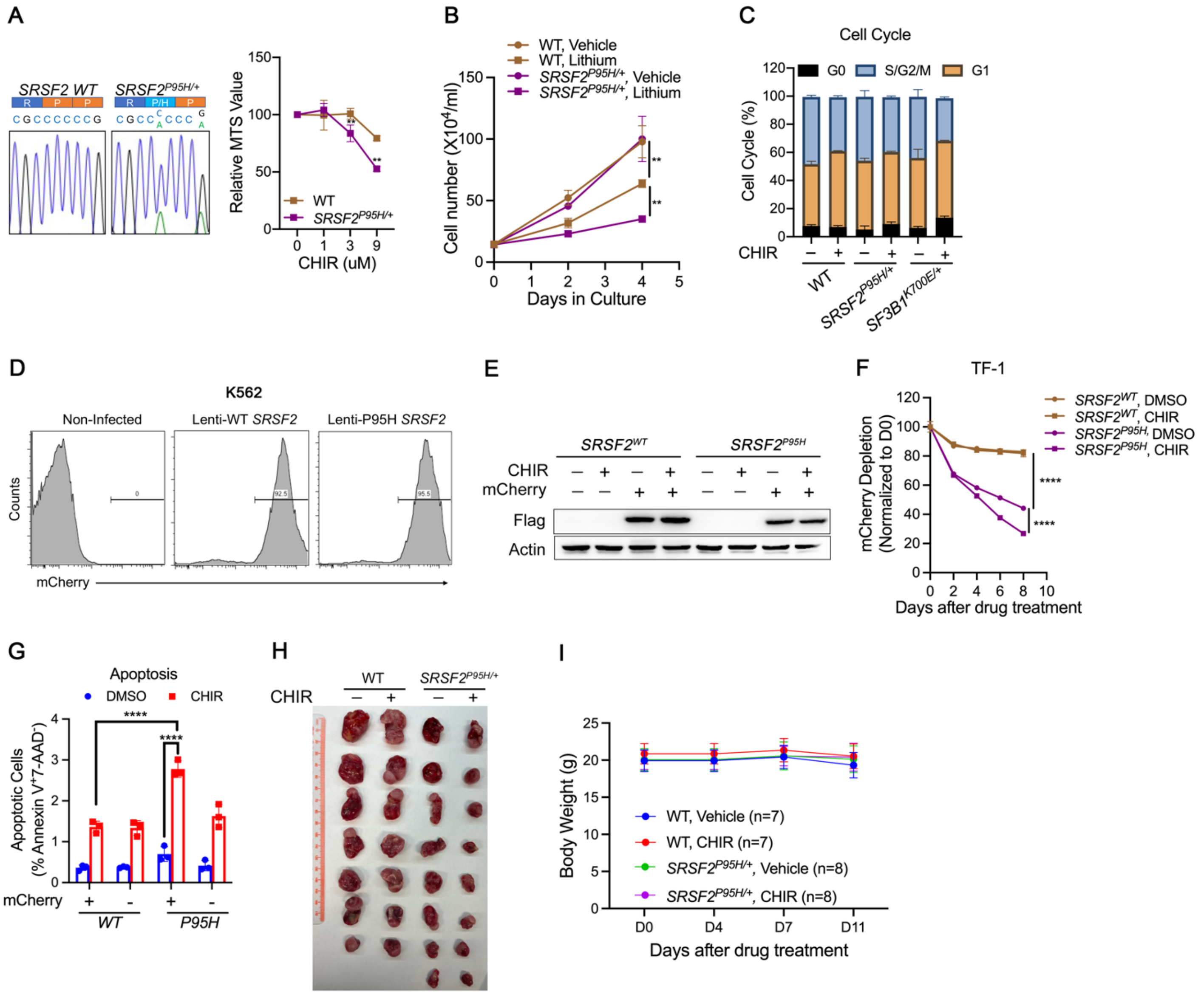


**Figure S1: supplement to Figure 1.** (A) The *SRSF2*$^{P95H/+}$ mutation was analyzed by Sanger sequencing of genomic DNA (gDNA) isolated from isogenic K562 *WT* and *SRSF2*$^{P95H/+}$ cells (left). *WT* and *SRSF2*$^{P95H/+}$ cells were treated with increasing concentrations of CHIR for 3 days. Cell numbers were then determined by MTS absorbance. Data are presented as mean ± SD. ** indicates $p$ < 0.01 (Student's $t$ test). (B) *WT* and *SRSF2*$^{P95H/+}$ cells were cultured with vehicle or 10mM Lithium, and cell numbers were counted at day 2 and day 4. (C) Percentage of cells in G0, S/G2/M, and G1 based on Ki67 and DAPI cytometric analysis of isogenic K562 *WT*, *SRSF2*$^{P95H/+}$ and *SF3B1*$^{K700E/+}$ cells treated with DMSO or 3μM CHIR *in vitro* for 4 d. There were no significant differences among the groups (two-tailed Chi-squared test). (D) Validation of lentiviral infection efficacy by flow cytometric analysis of mCherry$^{+}$ in K562 cells. (E)

Immunoblotting of flag-tag in sorted $mCherry^+$ TF-1 cells. (F) Percentage of $mCherry^+$ cells over 9 days in culture normalized to day 0. (G) Percentage of early-apoptotic cells based on flow cytometric analysis of 7-AAD and Annexin V in $\textit{SRSF2}^{wt}$- or $\textit{SRSF2}^{P95H}$-expressing TF-1 cells treated with DMSO or 1μM CHIR in vitro for 6 days. (H-I) Representative images of tumor (H), and mouse body weight (I) after 13 days of treatment in NSG mice subcutaneously implanted with K562 isogenic cells with or without $\textit{SRSF2}^{P95H/+}$ mutation. Data in B, F, and G are presented as the mean ± SD. ** $p < 0.01$, and **** $p < 0.0001$ (2-way ANOVA with Sidak's multiple comparisons test).

**Figure S2**

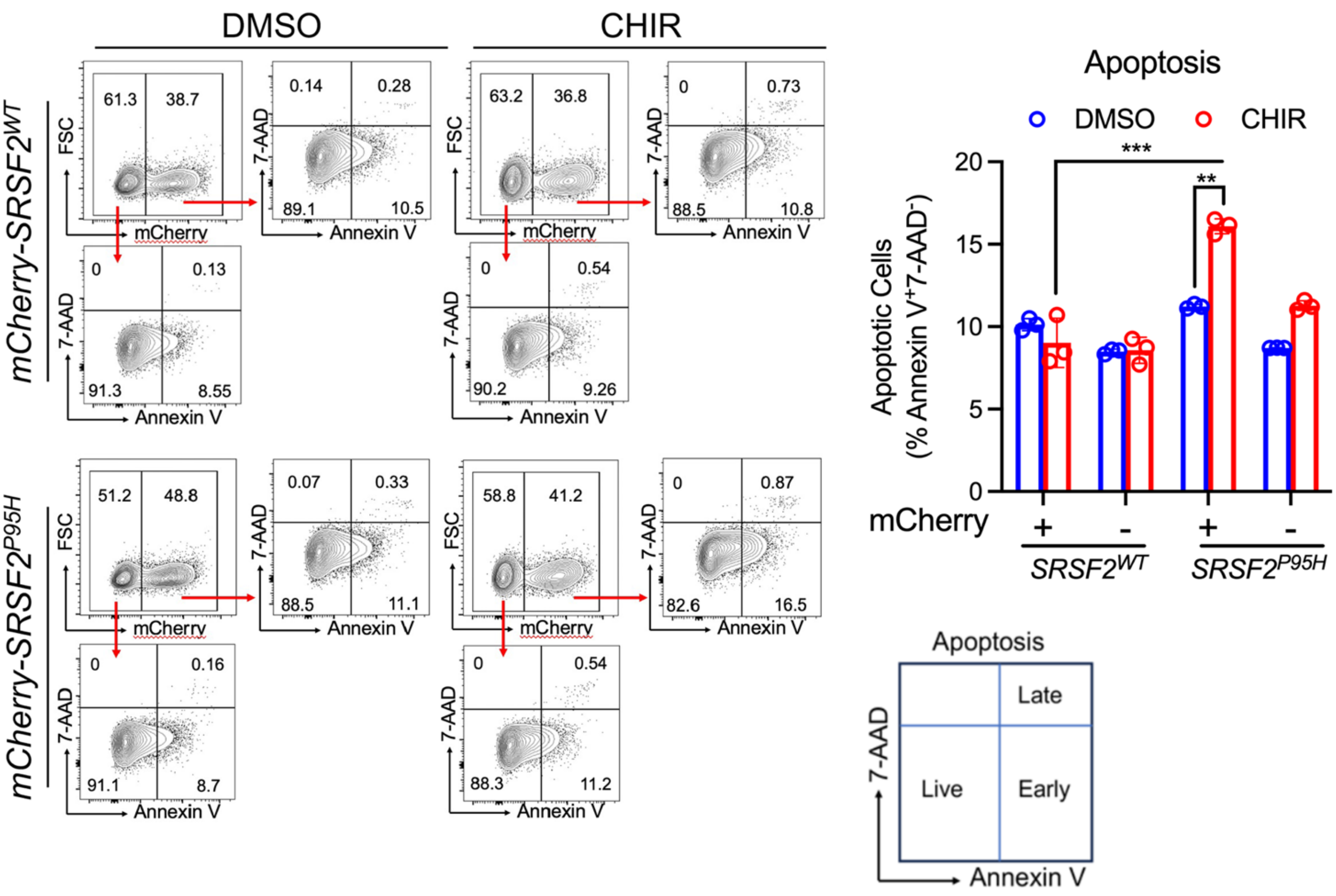


**Figure S2: supplement to Figure 2.** Representative flow cytometric analysis and quantification of apoptosis in primary cells from a patient with AML overexpressing either *SRSF2*$^{+}$ or *SRSF2*$^{P95H}$, as measured by Annexin V and 7-AAD staining in absence or presence of 3uM CHIR (similar to Figure 2D but with primary cells from a different AML patient. Data are presented as mean ± SD. ** $p < 0.01$ and *** $p < 0.001$ (2-way ANOVA with Sidak's multiple comparisons test).

### Figure S3

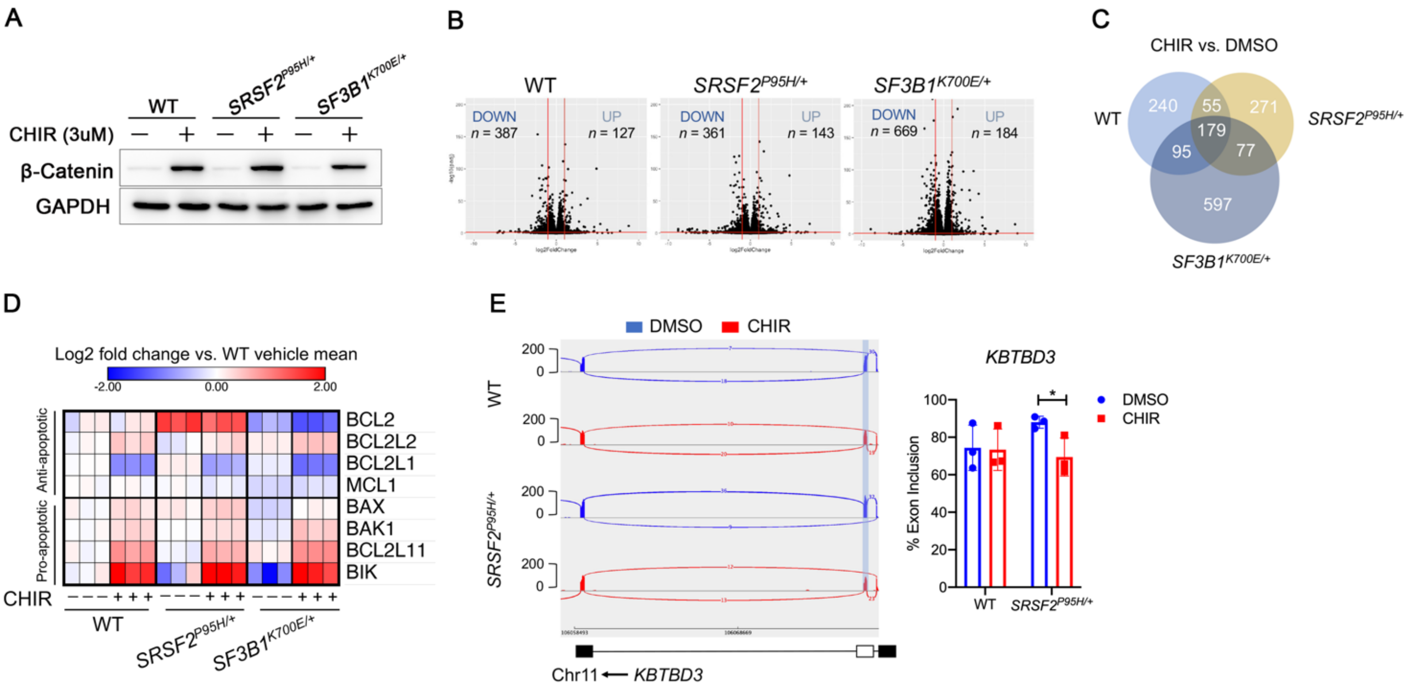


**Figure S3: supplement to Figure 3.** (A) Western blot analysis of β-catenin protein expression to validate GSK-3 inhibition by CHIR in K562 *WT*, *SRSF2*$^{P95H/+}$ and *SF3B1*$^{K700E/+}$ cells for RNA-seq. (B) Volcano plot analysis of all transcripts in CHIR versus DMSO treated *WT*, *SRSF2*$^{P95H/+}$ and *SF3B1*$^{K700E/+}$ K562 cells. Log2[FoldChange] was calculated for each transcript using normalized reads counts. (C) Venn diagram showing the numbers of overlapping differentially expressed genes between CHIR and DMSO treated *WT*, *SRSF2*$^{P95H/+}$ and *SF3B1*$^{K700E/+}$ K562 cells. (D) Heat map showing z-scored expression of *BCL2* family genes compared to mean expression for untreated parental cells. (E) Sashimi plots of *KBTBD3* (left) and quantification of exon inclusion percentage (right) in *WT* and *SRSF2*$^{P95H/+}$ cells treated with DMSO or CHIR. * $p$<0.05 (2-way ANOVA with Sidak's multiple comparisons test).

### Figure S4

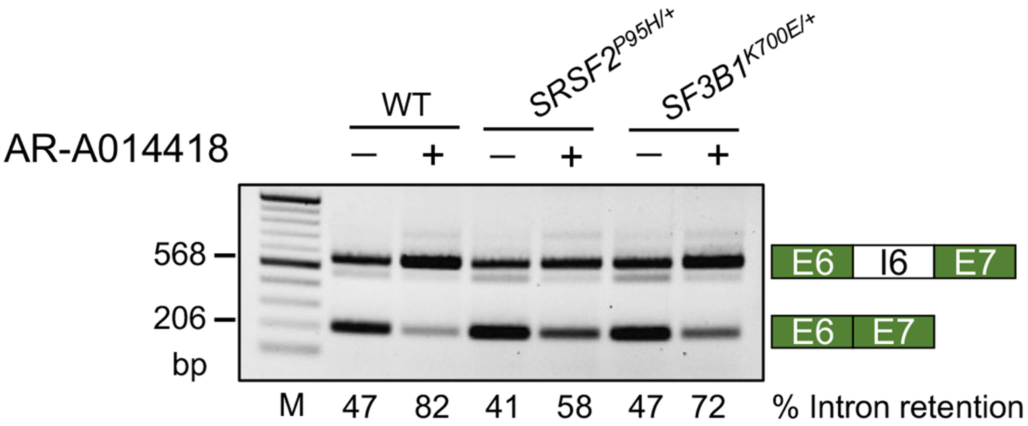


**Figure S4: supplement to Figure 4.** Representative RT-PCR with primers spanning exon 6 (E6), intron 6 (I6), and exon 7 (E7) of *PINK1* in *WT*, *SRSF2*$^{P95H/+}$, *SF3B1*$^{K700E/+}$ K562 cells treated with DMSO or 20uM AR-A014418 for 24h. M: DNA marker.

### Figure S5

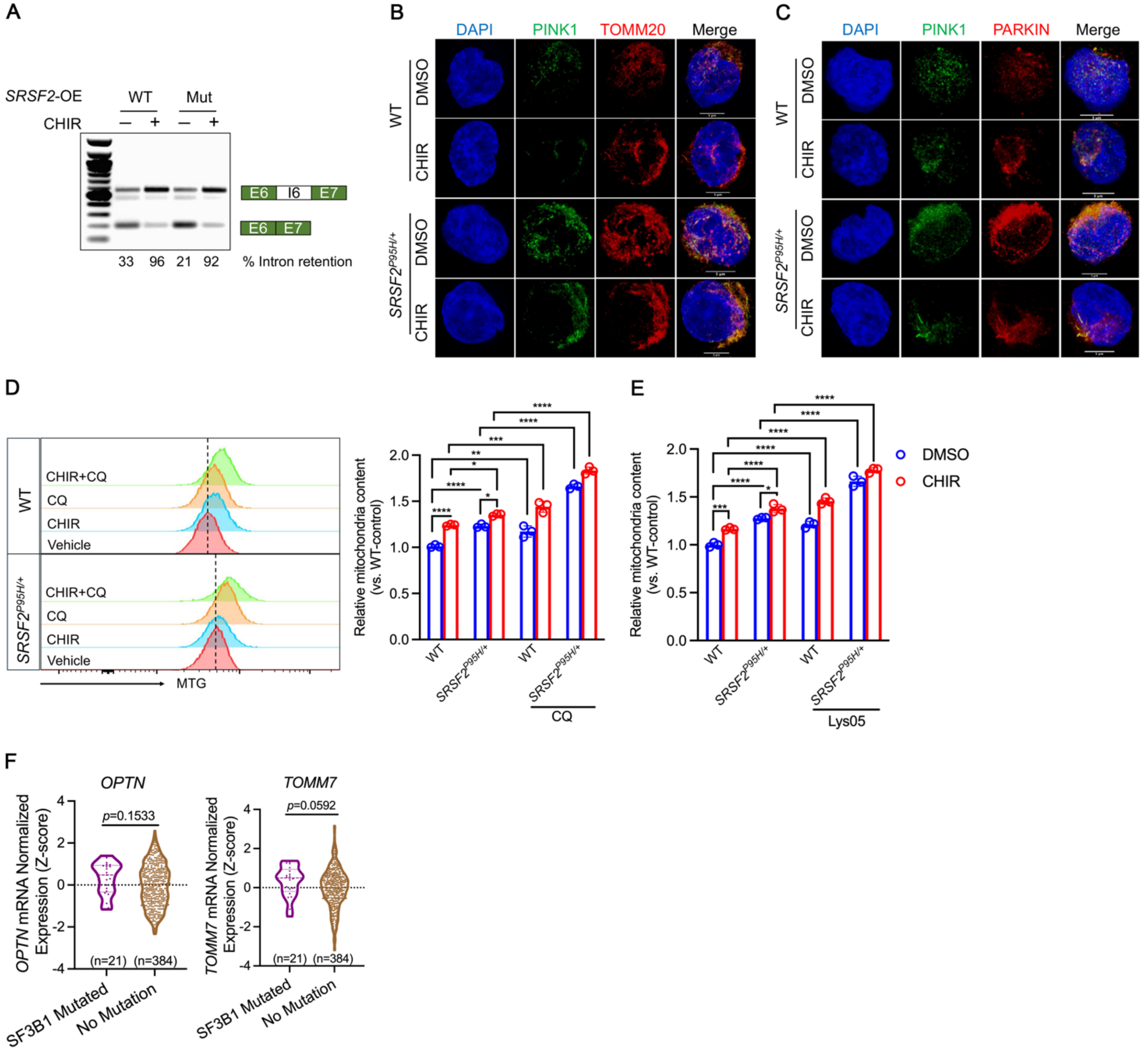


**Figure S5: supplement to Figure 5.** (A) Representative RT-PCR results of *PINK1* splicing in TF-1 cells expressing either *WT* or *P95H* mutant *SRSF2* treated with DMSO or 1μM CHIR for 24h. (B) Representative confocal images of mitochondria (red, TOMM20$^{+}$) and PINK1 (green) in K562 *WT* or *SRSF2$^{P95H/+}$* cells treated with DMSO or 3uM CHIR for 2d shown as 3D stack. Nuclei are stained with DAPI (blue). (C) Representative confocal images of PINK1 (green) and PARKIN (red) in K562 *WT* or *SRSF2$^{P95H/+}$* cells treated with DMSO or 3uM CHIR for 2d shown as 3D stack. Nuclei are stained with DAPI (blue). (D-E) Bar graph shows mitochondrial content determined by mito-tracker green (MTG) staining in *WT* and *SRSF2*$^{P95H/+}$ K562 treated with or without 2uM CHIR for 48h. Mitochondrial net flux was calculated by mitochondrial accumulation in the presence of 100uM Chloroquine (D) or 50uM Lys05 (E) for 4h. Data are presented as the mean ± SD. *$p$< 0.05, **$p$ < 0.01, ***$p$ < 0.001, ****$p$ < 0.0001 (2-way ANOVA with Sidak's multiple comparisons test). (F) Violin plot of *OPTN* and *TOMM7* normalized expression in AML patients in the TCGA dataset (n = 405) with or without mutations in *SF3B1*. Statistical analysis was performed using two-tailed Mann-Whitney test.

**Figure S6**

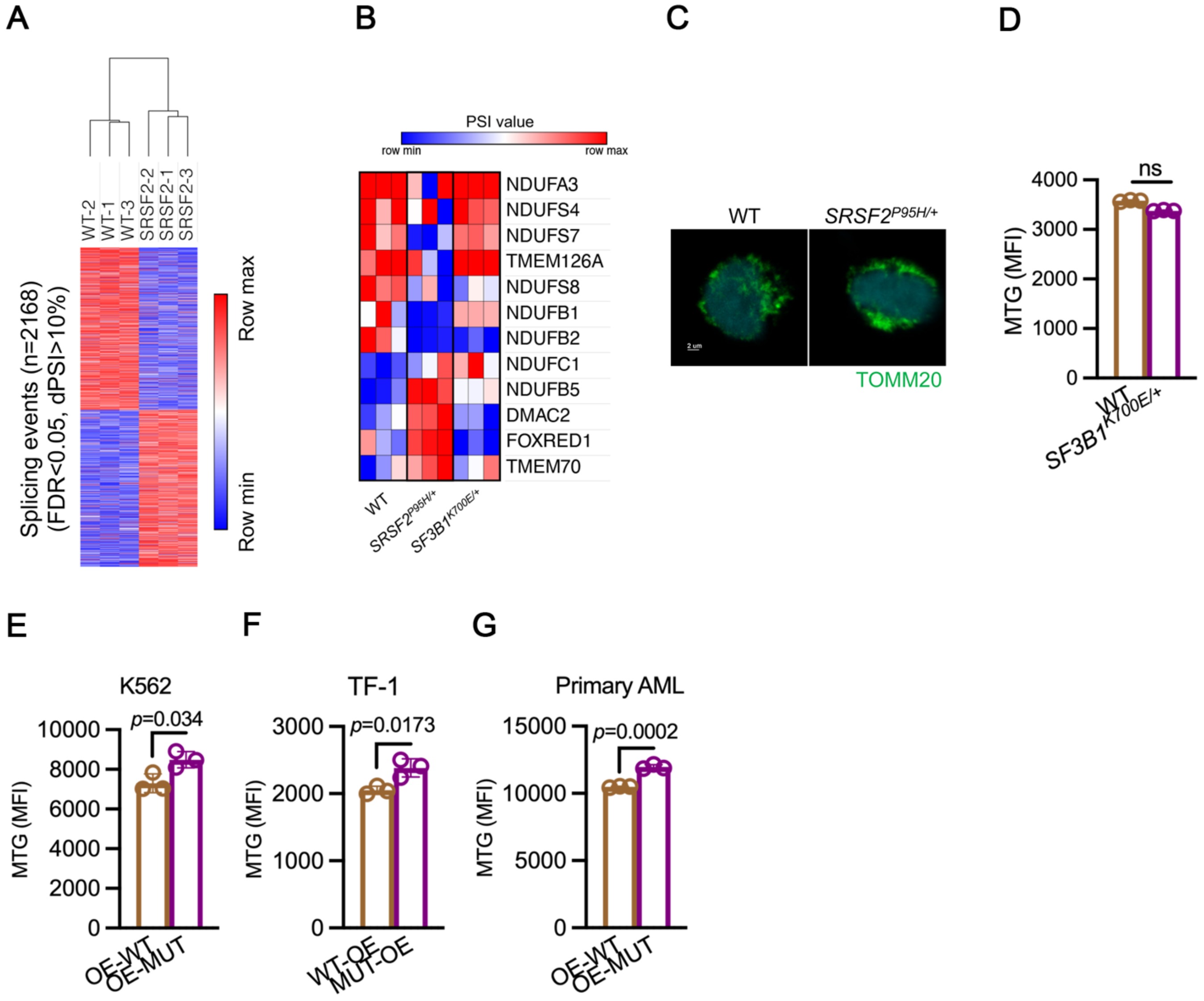


**Figure S6: supplement to Figure 6.** (A) Heat map showing splicing events using the 2168 differentially spliced isoforms between *WT* and *SRSF2*$^{P95H/+}$ cells (FDR < 0.05, dPSI > 10%). (B) Heat map showing splicing changes (PSI) for mRNAs encoding mitochondrial respiratory chain complex I proteins that are differentially spliced in *SRSF2*$^{P95H/+}$ cells compared to WT and *SF3B1*$^{K700E/+}$ cells. (C) Representative confocal images of mitochondria (TOMM20$^{+}$) in WT and *SRSF2*$^{P95H/+}$ cells are shown. (D) Quantification of mitochondrial mass by MTG staining in WT and *SF3B1*$^{K700E/+}$ cells. ns: not significant. (E-G) Quantification of mitochondrial mass by MTG staining in K562 (E), TF-1 (F), and primary AML (G) cells overexpressing *WT* or mutant (mut) *SRSF2*. Data are presented as mean ± SD. *p* values were determined by Student's *t* test (D-G).

### Figure S7

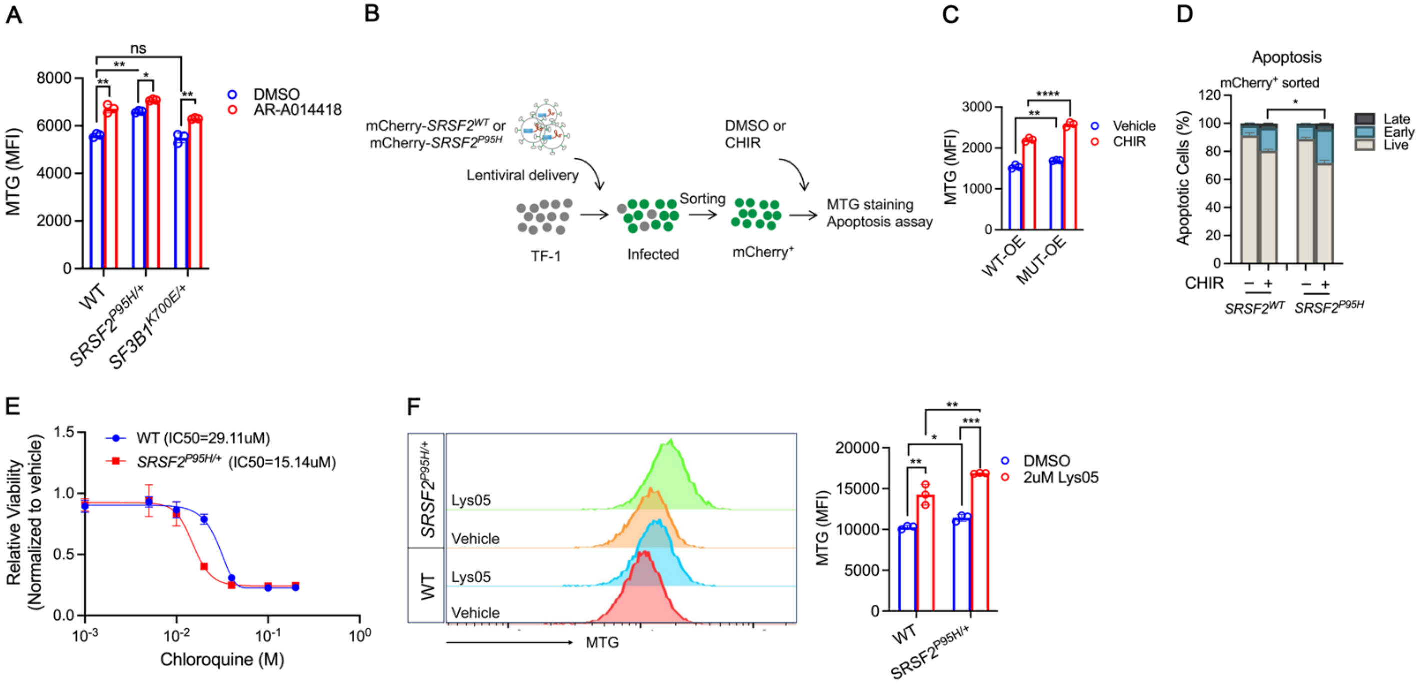


**Figure S7: supplement to Figure 7.** (A) Quantification of mitochondrial mass by MTG staining in WT, *SRSF2*$^{P95H/+}$, and *SF3B1*$^{K700E/+}$ cells treated with DMSO or 20μM AR-A014418 for 48h. (B) Schematic of lentivirus infection in TF-1 cells for MTG staining and apoptosis assay. (C) Quantification of mitochondrial mass by MTG staining and (D) apoptosis in TF-1 cells overexpressing *WT* or mutant (mut) *SRSF2* treated with DMSO or 1μM CHIR for 8d. (E) Chloroquine dose-response curves of *WT* and *SRSF2*$^{P95H/+}$ cells. (F) Quantification of mitochondrial mass by MTG staining in *WT* and *SRSF2*$^{P95H/+}$ cells treated with vehicle or 2uM Lys05 for 4d. Data in A, C, and F are presented as the mean ± SD. *$p < 0.05$, **$p < 0.01$, ***$p < 0.001$, and ****$p < 0.0001$ (2-way ANOVA with Sidak's multiple comparisons test). In D, * indicates $p < 0.05$ (two-tailed Chi-squared test).
